# Glutamine codon-driven translational readthrough reveals context-dependent stop codon decoding fidelity

**DOI:** 10.64898/2026.05.08.723929

**Authors:** Jessica M. Leslie, Kaitlin Morse, Sarah Swerdlow, Gloria A. Brar, Elçin Ünal

## Abstract

Deviations from the canonical genetic code include reassignment of UAA/UAG stop codons to glutamine in divergent eukaryotes, and tRNA^Gln^ has been shown to mediate near-cognate stop codon readthrough in canonical-code organisms. However, the sequence determinants and mechanistic basis of this decoding event remain poorly understood. Using ribosome profiling, quantitative immunoblotting, and mass spectrometry in *Saccharomyces cerevisiae*, we demonstrate that premature stop codon readthrough efficiency is governed by both local glutamine codon context and the global glutamine codon content of the mRNA. A QXQ motif flanking the stop codon promotes baseline readthrough, which is amplified in proportion to total transcript glutamine codon abundance. Mass spectrometry confirms that glutamine is specifically inserted at the premature stop, with no flanking miscoding, implicating tRNA^Gln^ competition with the release factor as the mechanistic basis of readthrough. Consistent with this model, yeast proteins terminating in short C-terminal glutamine repeats are evolutionarily enriched for strong stop codon contexts, suggesting selective pressure to reinforce termination fidelity at readthrough-prone loci.

## INTRODUCTION

Accurate decoding of the genetic code is fundamental to cellular life. In eukaryotes, translation termination is orchestrated by the release factors eRF1 and eRF3: eRF1 recognizes the stop codon in the ribosomal A site and recruits eRF3, whose GTPase activity triggers polypeptide release and ribosome recycling (1–3). Although termination is generally highly efficient, both *cis*-acting sequence features and *trans*-acting factors can promote stop codon readthrough, whereby an amino acid is erroneously incorporated in place of the stop codon, with important consequences for gene expression and proteome integrity (3–6).

Termination efficiency reflects the combined influence of local sequence context and *trans*-acting regulators. Transcriptome-wide analyses in yeast and mammalian cells demonstrate a hierarchy for stop codon strength (TAA>TAG>TGA, where TGA is most prone to readthrough), and support structural and biochemical data indicating that that termination is further enhanced when a purine (A/G) occupies the +4 position immediately downstream of the stop codon (7–9). The poly(A) tail and its binding protein PABP/Pab1 also enhance stop codon fidelity by engaging eRF3 (10, 11), and mRNA secondary structures downstream of the stop codon can suppress efficient termination in certain viral and metazoan transcripts (12–15). Among the *cis*-acting elements known to promote readthrough, the QXQ motif, in which the stop codon is flanked on both sides by glutamine codons, has been identified as a high-readthrough context in studies from yeast and mammalian cells (16–21). Peptide sequencing of readthrough products at QXQ sites reveals a mixture of incorporated amino acids, implicating near-cognate tRNA mispairing at the first or third codon position (19, 20). How glutamine codons mechanistically influence decoding fidelity at the stop codon, and whether sequence features beyond the immediate stop codon context contribute to readthrough efficiency, remains unclear.

Although the genetic code is considered universal, exceptions exist across bacteria and eukaryotes (22, 23). The most frequently observed eukaryotic deviation is stop codon reassignment. Among eukaryotic stop codon reassignments, the most common deviation, in which TAA and TAG are read as glutamine through the evolution of a tRNA^Gln^ with complementary anticodon identity, has independently evolved in diverse unicellular eukaryotes, including ciliates, green algae, oxymonads, and diplomonads (24–28). This recurrent evolutionary adaptation highlights tRNA^Gln^ as uniquely predisposed to acquire stop codon-decoding capacity, likely reflecting intrinsic near-cognate pairing potential at TAA and TAG codons that persists even in organisms retaining a canonical code.

Promoting readthrough at stop codons also has direct relevance to human health, as premature stop mutations account for at least 12% of all disease-causing mutations (29). Therapeutic strategies exploiting readthrough, including aminoglycoside antibiotics and engineered suppressor tRNAs, have attracted considerable interest (30–32). Deeper mechanistic understanding of how near cognate tRNAs compete with eRF1 at premature stop codons will therefore be important for refining these approaches.

Here, we uncover an unexpected role for polyQ tracts in promoting stop codon readthrough. We demonstrate that both local QXQ motifs and the broader glutamine codon composition of an mRNA contribute to readthrough efficiency through distinct mechanisms, that tRNA^Gln^ is specifically inserted at the premature stop codon without flanking loss of fidelity, and that nonsense mediated decay (NMD) factors negatively regulate this readthrough event. Finally, we find that yeast genes encoding C-terminal short glutamine repeats show a bias toward strong stop codon contexts, consistent with selection to maintain termination fidelity at loci inherently prone to readthrough. Together, our findings establish a framework for how glutamine codon usage shapes stop codon decoding fidelity and provide new insight into the evolutionary and mechanistic basis of tRNA^Gln^–mediated readthrough.

## RESULTS

### Nonsense mutations within the QLC domain of Snf5 result in unusually elevated levels of translational readthrough

A prior genetic screen in our laboratory uncovered mutations in several subunits of the SWI/SNF complex, revealing an unexpected role for this conserved chromatin remodeler in transcriptional repression (33). Particularly relevant to the current study were two nonsense mutations in *SNF5, snf5-Q225X* and *snf5-Q267X* (Figure S1A). Snf5 is a regulatory subunit that recruits SWI/SNF to promoter regions via interactions between its N-terminal glutamine-rich low-complexity (QLC) domain and acidic transcription factors (34). Notably, both mutations fall within this QLC domain.

We initially focused on *snf5-Q225X,* which is predicted to generate a truncated protein lacking ∼75% of Snf5, including a portion of the QLC domain (Figure S1A). Unexpectedly, *snf5-Q225X* mutants exhibited wild-type (WT)-like growth by colony size and doubling time and did not display the severe growth defects characteristic of the *snf5Δ* null mutant (Figures 1A–B; Figure S1B).

**Figure 1.**
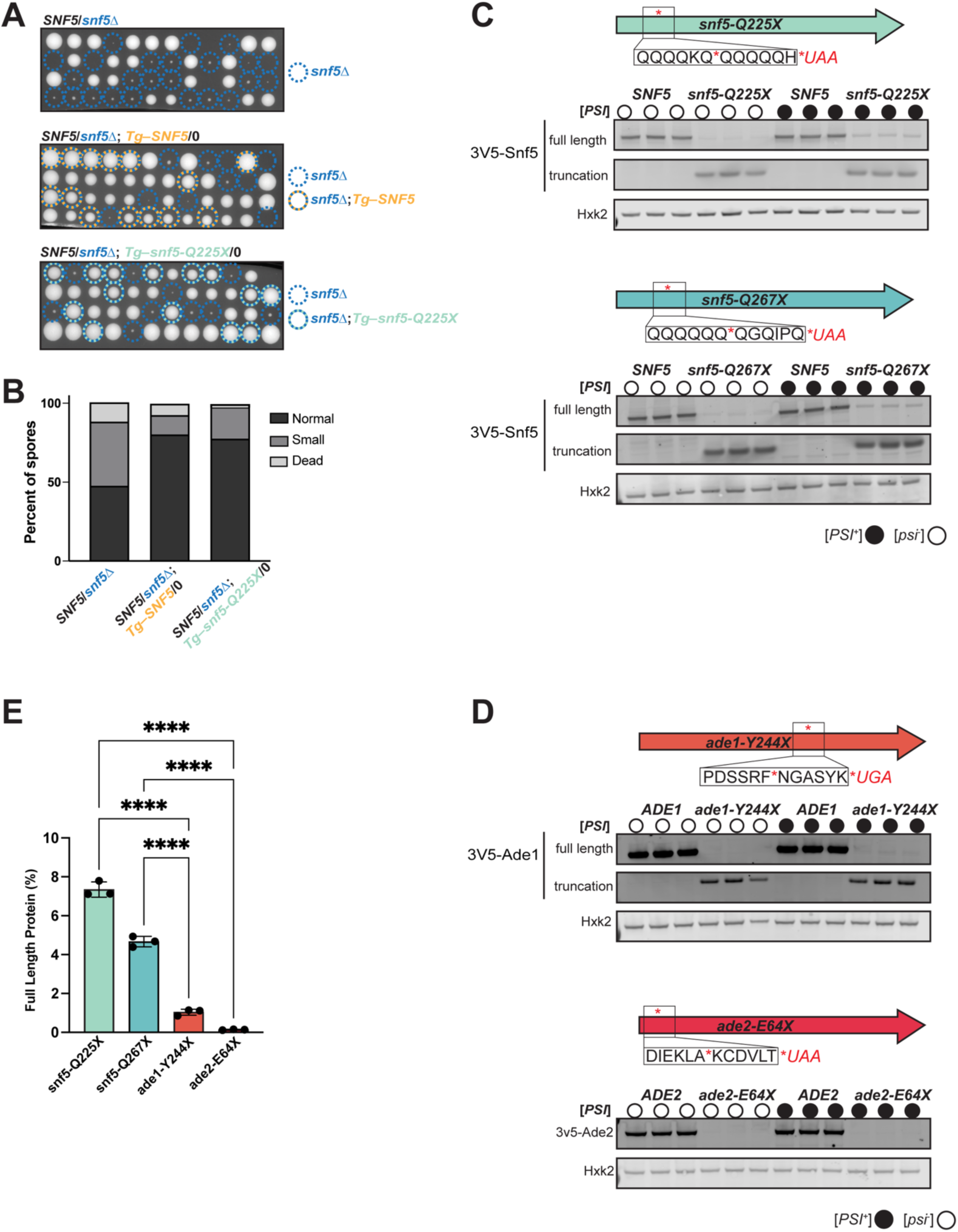
Nonsense mutations within the QLC domain of Snf5 result in unusually elevated levels of translational readthrough. (A) Tetrad dissections of hemizygous strains: snf5△ (top), SNF5 rescue (middle) and *snf5Q225X* (bottom). (B) Quantification of normal sized, small and dead spores in for each dissection in (A). (C) Semi-quantitative immunoblots for Snf5 (3v5-*snf5Q225X* and 3v5-*snf5Q267X*) premature stop alleles, [*PSI+*]/[*psi-*] as indicated. (D) Representative semi-quantitative immunoblots for Ade1 (3v5-*ade1Y244X*) and Ade2 (3v5-*ade2E64X*), [*PSI+*]/[*psi-*] as indicated. (E) Quantification of premature stop [*PSI+*] normalized to full length [*PSI+*] for each protein shown in (C) and (D). Three biological replicates are plotted, and each represents the average of two technical replicates. Statistical significance was determined by a one-way analysis of variance (ANOVA) adjusted for multiple comparisons with Tukey’s multiple comparisons test (****p_adj_<0.0001).

To determine whether the N-terminal 225 amino acids of Snf5 are sufficient for function, we constructed a clean truncation allele (*snf5^trunc^*) by deleting all coding sequences downstream of the Q225 position. This allele recapitulated the slow-growth phenotype of *snf5Δ* cells (Figure S1C), indicating that the truncated protein alone cannot support WT growth. We further ruled out reversion or second-site suppression as explanations for the near-WT growth of the *snf5-Q225X* strains: sequencing revealed no additional mutations, WT-like growth was observed in fresh segregants following outcrossing, and *de novo* introduction of *snf5-Q225X* into an independent background reproduced the original phenotype ((33); Figure 1A). We therefore examined whether translational readthrough of the premature stop codon permitted expression of full-length Snf5. Immunoblotting of 3xV5-tagged *snf5-Q225X* strains revealed that the mutant allele produced ∼7% of full-length Snf5 relative to WT (Figures 1C and 1E). Similarly, *snf5-Q267X* yielded ∼5% full-length protein (Figures 1C and 1E). Thus, both alleles undergo appreciable stop codon readthrough, generating sufficient Snf5 to support normal growth.

The genetic screen and subsequent experiments described above were conducted in the W303 strain background, which harbors the [*PSI*⁺] prion, an amyloid form of the translation termination factor Sup35 (eRF3) that promotes readthrough by sequestering functional eRF3 into aggregates (Figure S1G; (35)). [*PSI*^+^] presence and strength are traditionally assessed by a qualitative colony color assay exploiting nonsense alleles of adenine biosynthesis genes, *ade1-14* and *ade2-1*, which encode enzymes whose loss causes accumulation of a red pigment (36). To determine whether [*PSI*^+^] contributes to *snf5-Q225X* readthrough, we obtained [*PSI*^+^] and [*psi*^-^] W303 strains and confirmed their prion status using a Sup35NM-GFP reporter, which forms cytoplasmic puncta in [*PSI*⁺] strains but remains diffuse in [*psi*⁻] cells (Figure S1H; (37, 38)). Immunoblotting revealed that readthrough at *snf5-Q225X* was significantly enhanced in [*PSI*⁺] strains but remained detectable in [*psi*⁻] backgrounds (Figure S1F), indicating that basal readthrough occurs independently of prion status.

Although the colony color assay has long served as a qualitative proxy for readthrough, quantitative comparisons of protein output across nonsense alleles have been limited. To determine whether the Snf5 readthrough levels we observed were unusually high, we generated N-terminal 3xV5-tagged versions of the canonical readthrough reporters *ade1-14* (*ade1-Y244X*) and *ade2-1* (*ade2-E64X*) in [*PSI*⁺] and [*psi*⁻] backgrounds and compared full-length protein levels to WT controls by quantitative immunoblotting (Figure 1D). Remarkably, *snf5-Q225X* produced ∼56-fold more full-length protein than *ade2-1* and ∼8-fold more than *ade1-14,* while *snf5-Q267X* produced ∼35-fold more full-length protein than *ade2-1* and ∼4.5-fold more than *ade1-14*. On average, *snf5-Q225X*, *ade1-Y244X* and *ade2-E64X* transcripts were 88%, 45% and 24% as abundant as their WT counterparts (Figure S1E). Although transcript levels varied 2-4-fold among alleles, readthrough levels differed by as much as 50-fold, indicating that differences in mRNA abundance alone cannot account for the magnitude of the readthrough effect. Together, these results reveal a striking disparity in stop codon readthrough between *SNF5* and the canonical auxotrophic alleles, underscoring gene- and context-dependent variability in translational termination fidelity.

### Premature stop codon readthrough is partially determined by local glutamine and stop codon context

The high-readthrough *SNF5* alleles share two sequence features that have each been implicated in promoting stop codon readthrough. The first is a cytosine at the +4 position immediately downstream of the stop codon, which has been suggested to reduce the efficiency of translational termination through reduced stacking interactions at the release factor site (9). The second is a QXQ motif flanking the stop codon, which has been shown to promote readthrough (19, 20). To determine whether these local sequence features account for the elevated readthrough we observed, we systematically altered the codon context surrounding the stop codon in the low-readthrough *ADE2* reporter, generating variants that test stop codon identity, the influence of 5′ or 3′ glutamine codons individually, or their combined effect (Figure 2A). Full-length Ade2 protein levels were then measured as a quantitative proxy for readthrough efficiency (Figure 2B–C).

**Figure 2.**
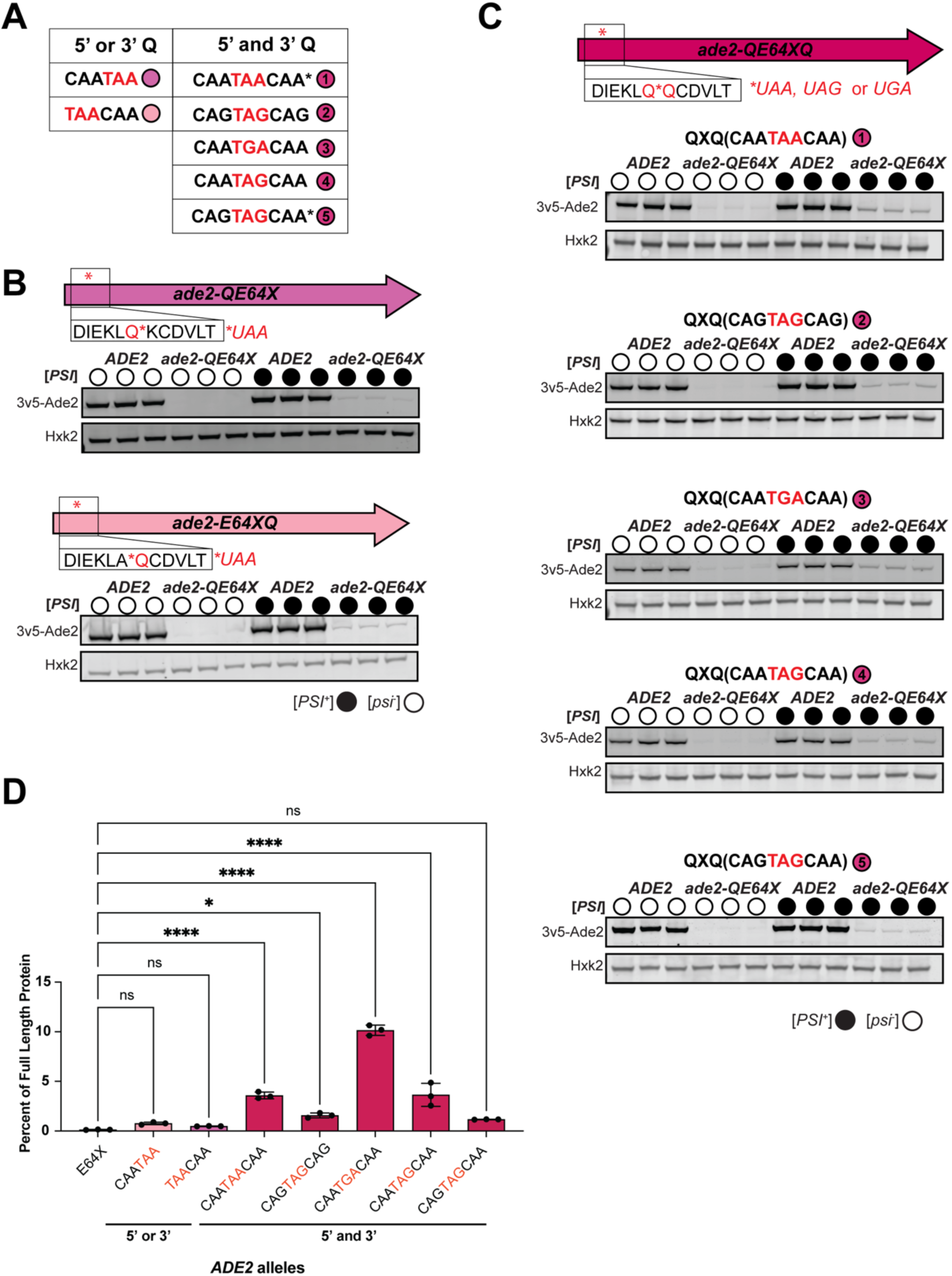
Premature stop codon readthrough is partially determined by local glutamine and stop codon context. (A) Table of Q and stop mutations generated in Ade2 at E64. (B) Representative semi-quantitative immunoblots for Ade2, 5’ (3v5-*ade2**Q**E64X,* CAA-TAA) or 3’ (3v5-*ade2E64X**Q**,* TAA-CAA) glutamines with premature stop alleles, [*PSI+*]/[*psi-*] as indicated. (C) Representative semi-quantitative immunoblots for Ade2, 3’ and 5’ glutamine premature stop alleles: CAA-TAA-CAA, CAG-TAG-CAG, CAA-TGA-CAA, CAA-TAG-CAA and CAG-TAG-CAA, [*PSI+*]/[*psi-*] as indicated. (D) Quantification of premature stop [*PSI+*] normalized to full length [*PSI+*] for each Ade2 variant shown in (B) and (C). Three biological replicates are plotted, and each represents the average of two technical replicates. Statistical significance was determined by a one-way analysis of variance (ANOVA) adjusted for multiple comparisons with Dunnett’s multiple comparisons test (*p_adj_ <0.05, *** p_adj_<0.001, ****p_adj_<0.0001).

Readthrough was highest when the stop codon was flanked on both sides by the CAA glutamine codons (Figure 2D). Among the three stop codons tested, UGA was most permissive, yielding ∼8% of full-length Ade2 protein, while UAG and UAA were less so, at ∼3% and ∼3.5%, respectively (Figure 2D). A small but statistically significant increase in readthrough was also observed when CAG codons flanked UAG and UAA stop codons, indicating that both the 5’ and 3’ glutamine codons independently contribute to readthrough (Figure 2D).

Notably, *ADE2* QXQ variants engineered to match the stop-proximal sequences of the *SNF5* alleles still displayed markedly lower readthrough than the corresponding *SNF5* alleles. For example, CAA-TAA-CAA context yielded 3.6% of full-length protein in Ade2 versus 7.4% in *snf5-Q225X*, and CAG-TAG-CAA yielded 1.6% in Ade2 versus 4.7% in *snf5-Q267X* (Figure 1E vs Figure 2D). Furthermore, substituting the more permissive CAA codon either 5′ or 3′ of the stop alone produced only modest, non-significant increases in readthrough, indicating that the combined flanking context is required for the full effect (Figure 2D). Codon modifications did not affect *ADE2* mRNA levels or Ade2 protein stability, as assessed by quantitative PCR and by protein-level measurements following addition of the proteasome inhibitor MG132, respectively (Figures S2C–D). Taken together, while flanking glutamine codons promote readthrough in a stop-codon-dependent manner; this local sequence effect is insufficient to explain the substantially elevated readthrough observed at premature stop codons within *SNF5*.

### Premature stop codon readthrough correlates with regional glutamine context

To identify additional sequence features that render *SNF5* uniquely permissive to readthrough, we examined differences in amino acid composition between *SNF5, ADE1* and *ADE2.* Among the three proteins, the frequency of each amino acid across all three proteins generally fell between 0.18-9.8%, except for glutamine (Q) usage in *SNF5*, which was a clear outlier at 16.7% Q (Figure 3A). Comparatively, *ADE1* and *ADE2* are relatively glutamine-depleted at 2.9% Q and 2.6% Q. These observations motivated us to ask whether other glutamine-enriched proteins are similarly permissive to premature stop codon readthrough.

**Figure 3.**
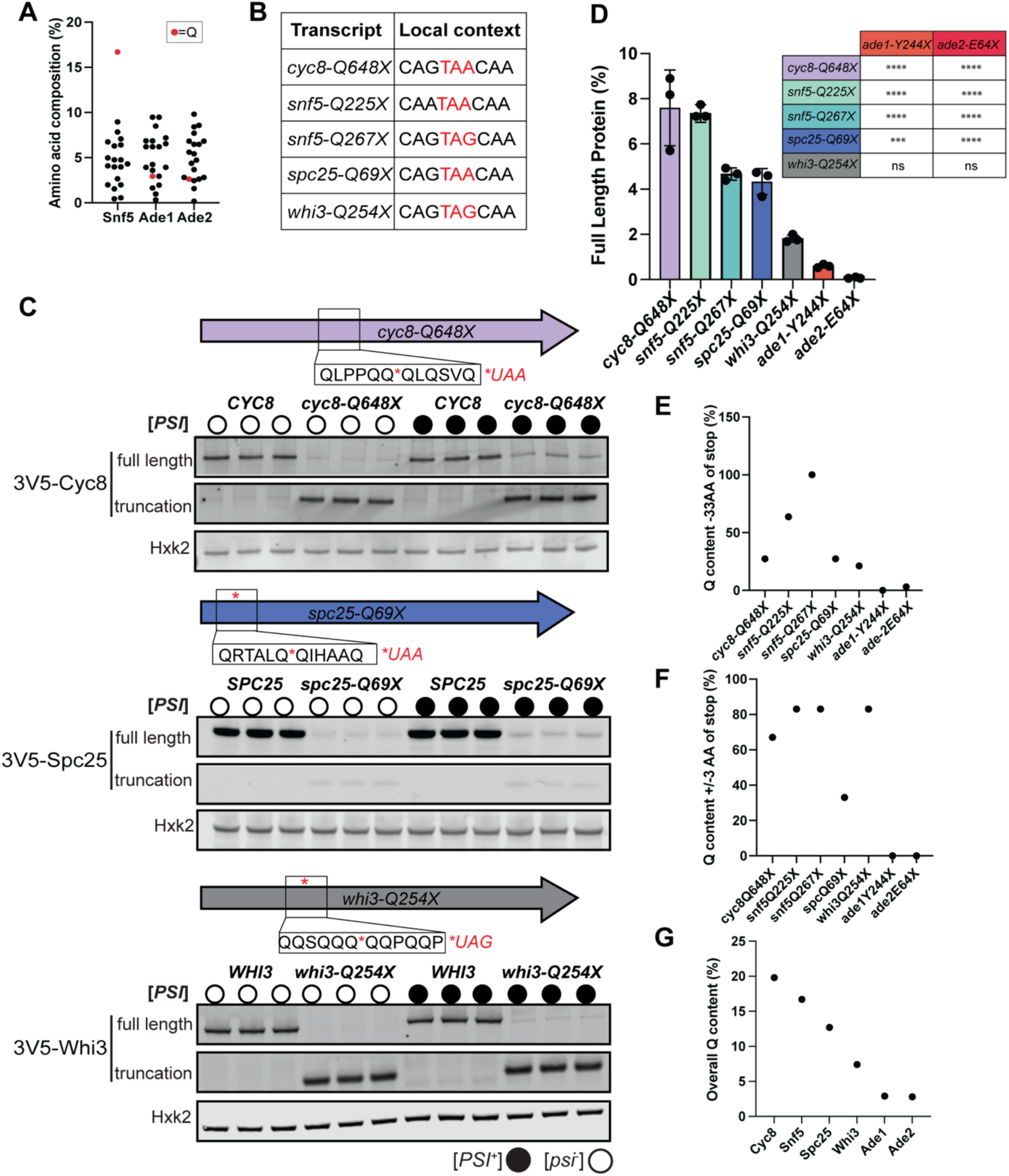
Premature stop codon readthrough correlates with regional glutamine context. (A) Percentages of each amino acid represented in Snf5, Ade1 and Ade2. Glutamine (Q) highlighted in red. (B) Table of Q and stop mutations represented in Q-rich alleles. (C) Representative semi-quantitative immunoblots for Cyc8, Spc25, and Whi3 premature stop alleles, [*PSI+*]/[*psi-*] as indicated. (D) Quantification of premature stop [*PSI+*] strains normalized to full length [*PSI+*] strains for each protein shown in C. Three biological replicates are plotted, and each represents the average of two technical replicates. Statistical significance was determined by a one-way analysis of variance (ANOVA) adjusted for multiple comparisons using Sidak’s multiple comparisons test (*** p_adj_<0.001, ****p_adj_<0.0001). (E) Glutamine percentage of the predicted ribosome exit tunnel length (∼33aa upstream of the premature stop) of all alleles in 3D. (F) Glutamine proportion of the 3 amino acids upstream and downstream (+/-) of the premature stop codon for all alleles in 3D. (G) Total glutamine percentage of candidate glutamine-rich (Cyc8, Snf5, Spc25, Whi3) proteins and non-glutamine rich controls (Ade1, Ade2).

We introduced 3xV5-tagged nonsense alleles into *WHI3, SPC25, and CYC8,* genes encoding glutamine-rich yeast proteins with Q content ranging from 7 to 20% (34). Each allele placed a UAG or UAA stop codon in a flanking glutamine codon context (CAG/CAA; Figure 3B). Compared to glutamine-depleted controls *ade2-E64X and ade2-Y244X*, these alleles yielded 2-8% full-length protein, with readthrough efficiency broadly correlating with total Q content of the encoded protein, but not with local Q context at the stop codon or glutamine codon density within the predicted ribosome exit tunnel (Figure 3E–G) (39). Although local glutamine and stop codon identity varied across the premature stop alleles tested, no consistent trends emerged from these local sequence features alone (Figure 3B). Taken together, these data suggest that the global abundance of glutamine codons within an mRNA contributes to readthrough efficiency beyond what is explained by local stop codon context alone. From here forward, we use “glutamine-rich” or “Q-rich” readthrough to distinguish this condition from isolated QXQ readthrough.

### Ribosome profiling confirms elevated stop codon readthrough in glutamine-rich transcripts

To directly quantify translation past premature stops, we performed ribosome profiling on three strain combinations harboring high-readthrough alleles: *cyc8-Q648X* [*PSI^+^*] and *snf5-Q225X* in both [*PSI^+^*] and [*psi^-^*] backgrounds (Figure 4A). All strains also carried *ade1-14* (*ade1-Y244X*) as an internal low-readthrough control. In-parallel immunoblotting confirmed that readthrough levels were consistent with prior experiments and colony growth assays validated [PSI] prion status in each strain (Figures S4A–D). We focused our analysis on 28-29nt ribosome-protected fragments, as most reads in this size range were biased towards a single reading frame (Figure S4F; see frame 1 for 28-29nts in all samples).

**Figure 4.**
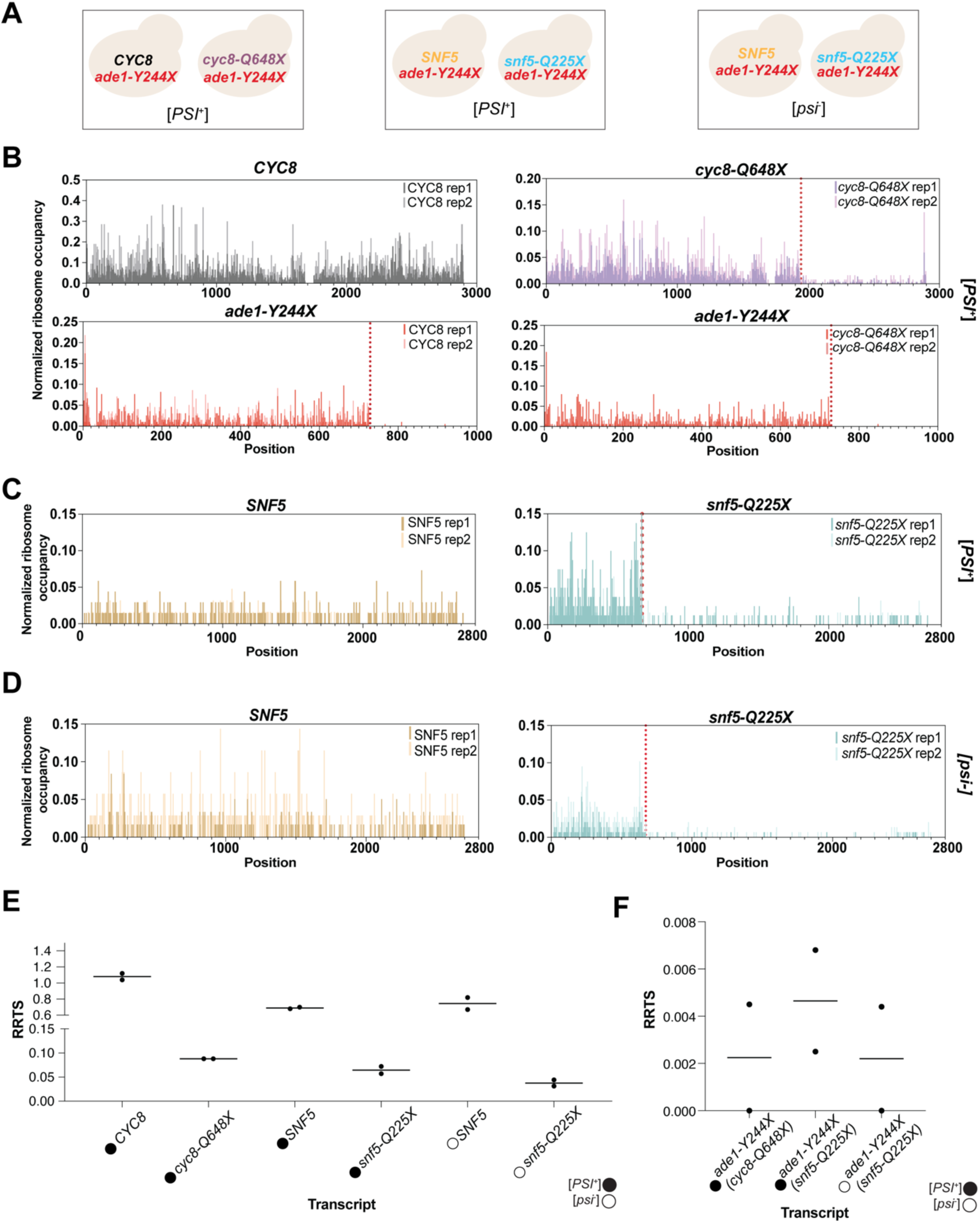
Ribosome profiling confirms elevated stop codon readthrough in glutamine-rich transcripts. (A) Schematic of strains tested with [*PSI*] status. (B) Normalized ribosome profiling traces of [*PSI+*] wildtype CYC8 (left) and *cyc8-Q648X* (right) with each strain’s corresponding *ade1-Y244X* control below. *Cyc8-Q648X* stop: 1944nt, *ade1-Y244X* stop: 729nt. Two replicates plotted. (C) Normalized ribosome profiling traces of [*PSI+*] wildtype SNF5 (left) and premature stop *snf5-Q225X* (right), *snf5-Q225X* stop: 672nt. Two replicates plotted. Corresponding *ade1-Y244X* control plots in S4I. (D) Normalized ribosome profiling traces of [*psi-*] wildtype SNF5 (left) and premature stop *snf5-Q225X* (right), *snf5-Q225X* stop: 672nt. Two replicates plotted. Corresponding *ade1-Y244X* control plots in S4J. (E) Ribosome readthrough scores (RRTSs) for *CYC8/cyc8-Q648X, SNF5/snf5-Q225X* [*PSI+*] and *SNF5/snf5-Q225X* [*psi-*] traces in (B)-(D), RRTS = 3’ ribosome densities post-stop/5’ ribosome densities pre-stop. [*PSI+*]/[*psi-*] as indicated. (F) Ribosome readthrough scores (RRTSs) for *ade1-Y244X* traces in 4B, S4I and S4J. RRTS = 3’ ribosome densities post-stop/5’ ribosome densities pre-stop.

We first confirmed readthrough at each locus by visualizing normalized ribosome footprint densities downstream of the premature stop codon (Figures 4B-D). Ribosome density past the premature stop was detectable in all conditions tested: *cyc8-Q648X* [*PSI+*], *snf5-Q225X* [*PSI+*] and *snf5-Q225X* [*psi-*]. Notably, *cyc8-Q648X* displayed ribosome density peaks near the canonical stop codon (Figure 4B), a pattern characteristic of ribosome pausing during termination and consistent with productive readthrough of the premature stop. To confirm that post premature stop translation proceeded in frame, we compared the proportion of first frame ribosome footprints upstream and downstream of the premature stop codon. Although a slight but reproducible decrease in the first frame occupancy was observed downstream of the premature stop in *cyc8-Q648X*, the majority of footprints in both *cyc8-Q648X* and *snf5-Q225X* remained in the first frame (Figure S4G-H), supporting the conclusion that ribosome profiling is capturing productive translational readthrough. An outlier frame distribution in replicate two of wild-type *SNF5* [*PSI+*] cells led us to individually investigate these replicates, and we concluded that replicate two shows a less biased distribution of frames (Figure S4E), suggesting that this replicate may have undergone incomplete nuclease digestion. We retained this replicate for downstream analysis as the overall trends remained consistent across replicates.

To compare readthrough levels quantitatively across transcripts, we calculated a Ribosome Readthrough Score (RRTS) by dividing normalized ribosome density in the 3’ region downstream of the premature stop by that in the 5’ region upstream, as previously described in Wangen *et al* (40). Both *cyc8-Q648X* and *snf5-Q225X* displayed higher RRTS values than the *ade1-Y244X* control, consistent with the immunoblot data (Figure 4E-F). In addition, *snf5-Q225X* readthrough was elevated in a [*PSI*⁺]-dependent manner (Figure 4E). Together, these results confirm that premature stop codon readthrough occurs at elevated levels in glutamine-rich transcripts.

### Glutamine-rich readthrough is largely refractory to canonical translational quality control pathways

To assess whether known translation-associated quality control pathways contribute to glutamine-rich readthrough beyond eRF3/Sup35, we tested candidate regulators of premature stop codon recognition and tRNA decoding fidelity, grouping them into positive regulators predicted to enhance readthrough (Itt1, Rps30b) and negative regulators predicted to suppress it (RQC, NMD).

We first examined the two candidate positive regulators. The E3 ubiquitin ligase Itt1, the yeast homolog of mammalian RNF14, promotes degradation of eRF1 stalled at stop codons (41, 42), and we therefore asked whether eRF1 turnover enhances readthrough at glutamine-rich premature stops. Deletion of *ITT1* had no effect on *snf5-Q225X* readthrough, indicating that eRF1 turnover does not significantly influence readthrough efficiency at glutamine-rich premature stops (Table 1; Table 1-Supplementary Data 1A). We also investigated the small ribosomal subunit protein Rps30b, which has recently been shown to promote near-cognate stop codon readthrough by tRNA^Gln^ through direct interaction with the tRNA^Gln^_CUG_ isoacceptor (43). We obtained strains with a point mutation (R10A) in Rps30b that specifically disrupts this interaction and introduced them into the *snf5-Q267X* background, as a *SNF5* allele harboring a TAG premature stop more closely matched to the tRNA^Gln^_CUG_ isoacceptor. Readthrough efficiency was unchanged in Rps30b-R10A strains relative to wild type, indicating that the Rps30b-tRNA^Gln^ interaction does not positively regulate glutamine-mediated stop codon bypass (Table 1; Table 1-Supplementary Data 1B).

**Table 1.**
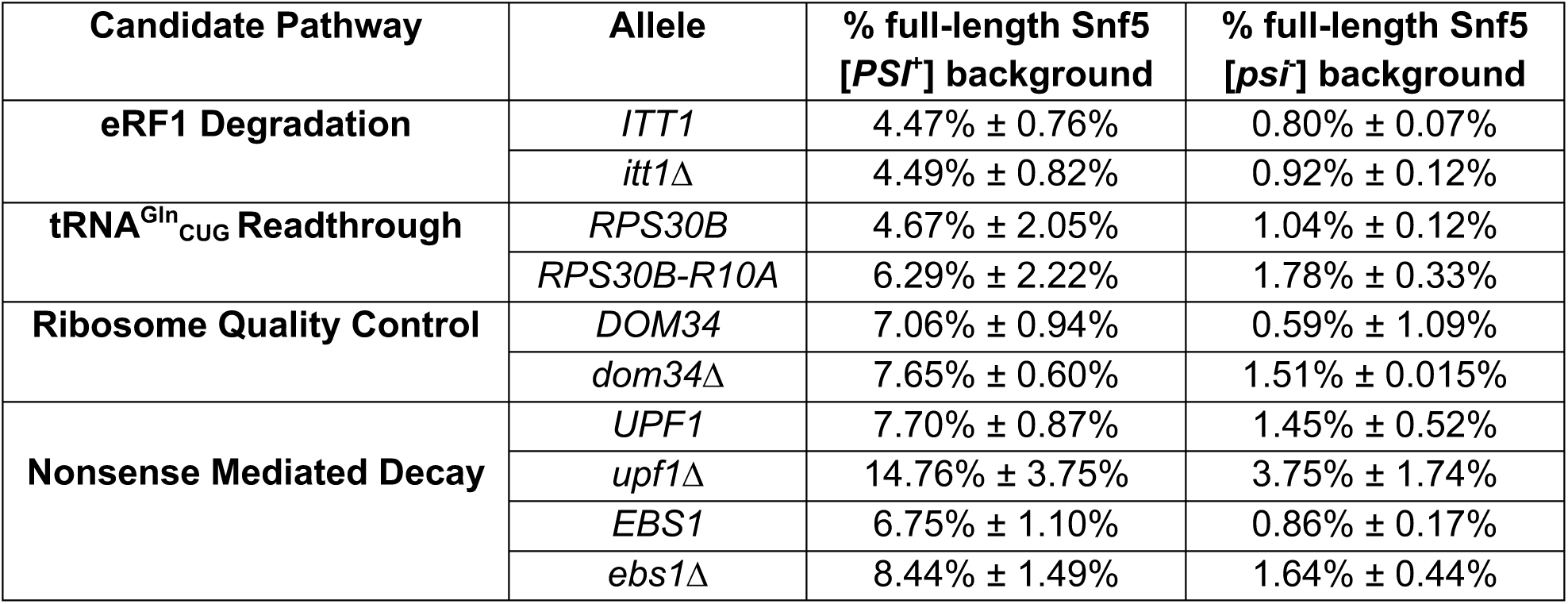
Results of testing candidate translational pathways for their involvement in snf5Q225X readthrough reported by immunoblot measurements (Figure S5A-D) percent of full-length protein in corresponding genotype). Values are reported as mean ± standard deviation (n=2 biological replicates, each measured in duplicate).

| Candidate Pathway | Allele | % full-length Snf5 [PSI<sup>+</sup>] background | % full-length Snf5 [psi<sup>-</sup>] background |
| --- | --- | --- | --- |
| **eRF1 Degradation** | *ITT1* | 4.47% ± 0.76% | 0.80% ± 0.07% |
|  | *itt1Δ* | 4.49% ± 0.82% | 0.92% ± 0.12% |
| **tRNA<sup>Gln</sup><sub>CUG</sub> Readthrough** | *RPS30B* | 4.67% ± 2.05% | 1.04% ± 0.12% |
|  | *RPS30B-R10A* | 6.29% ± 2.22% | 1.78% ± 0.33% |
| **Ribosome Quality Control** | *DOM34* | 7.06% ± 0.94% | 0.59% ± 1.09% |
|  | *dom34Δ* | 7.65% ± 0.60% | 1.51% ± 0.015% |
| **Nonsense Mediated Decay** | *UPF1* | 7.70% ± 0.87% | 1.45% ± 0.52% |
|  | *upf1Δ* | 14.76% ± 3.75% | 3.75% ± 1.74% |
|  | *EBS1* | 6.75% ± 1.10% | 0.86% ± 0.17% |
|  | *ebs1Δ* | 8.44% ± 1.49% | 1.64% ± 0.44% |

We next asked whether the ribosome-associated quality control (RQC) pathway, which detects and resolves stalled ribosomes at problematic sequences including polyA tracts and CGA codon repeats (44, 45), negatively regulates Q-rich readthrough. Loss of Dom34, which acts with Hbs1 to detect stalled ribosomes and promote their recycling (43), had no measurable effect on *snf5-Q225X* readthrough, indicating that canonical RQC surveillance does not contribute to suppression of polyQ-dependent readthrough (Table 1; Table 1-Supplementary Data 1C).

In contrast, examination of the nonsense-mediated decay (NMD) factors revealed a negative regulatory role. NMD targets premature termination codons through the RNA-dependent helicase/ATPase Upf1; mRNA degradation is initiated when downstream effectors, including Nmd4 and Ebs1, are recruited to the Upf1-Upf2-Upf3 complex to engage decapping and exonuclease activities (46). Deletion of *UPF1* or downstream effector *EBS1* increased *snf5-Q225X* readthrough, indicating that NMD factors act as negative regulators (Table 1; Table 1-Supplementary Data 1D). Importantly, *snf5-Q225X* transcript levels were largely unchanged in *upf1Δ* cells (Table 1-Supplementary Data 1E), indicating that this effect reflects a translational role for NMD factors in suppressing readthrough independently of their canonical function in mRNA degradation.

Taken together, these results demonstrate that the elevated readthrough observed at glutamine-rich premature stops in *SNF5* is negatively regulated by NMD factors, yet not otherwise modulated by canonical trans-acting quality control pathways, including RQC, ribosomal protein-tRNA^Gln^ interactions, or termination factor degradation.

### Glutamine is incorporated at the site of premature stop codon readthrough in Q-rich sequences

Given our observations that glutamine-rich sequences drive increased stop codon readthrough, we considered two models of near-cognate decoding (Figure 5A): (1) during translation of glutamine-rich sequences, tRNA^Gln^ is locally enriched and outcompetes the release factor at the stop codon, resulting in specific glutamine insertion; or (2) tRNA^Gln^ becomes limiting during synthesis of glutamine-rich proteins, causing a generalized loss of decoding fidelity and random miscoding at and around the stop codon, a model motivated by prior *in vitro* data (47). To distinguish between these possibilities, we overexpressed *cyc8-Q648X*, the allele displaying the highest level of readthrough among those tested, using a tetracycline-inducible system (48), and purified the readthrough product for mass spectrometry analysis (Figure S5A-B). Peptide mapping across two biological replicates (318 and 208 peptides, covering 74% and 59% of the protein sequence, respectively) revealed consistent and specific substitution of glutamine at the premature stop codon position (Figures 5B-C; Figures S5C). No miscoding was detected at flanking positions, and genomic sequencing confirmed that the nonsense mutation was intact in the expressed allele (Figure S5C). We therefore conclude that tRNA^Gln^ directly competes with the release factor at the premature stop codon, resulting in specific glutamine insertion rather than a general loss of decoding fidelity.

**Figure 5.**
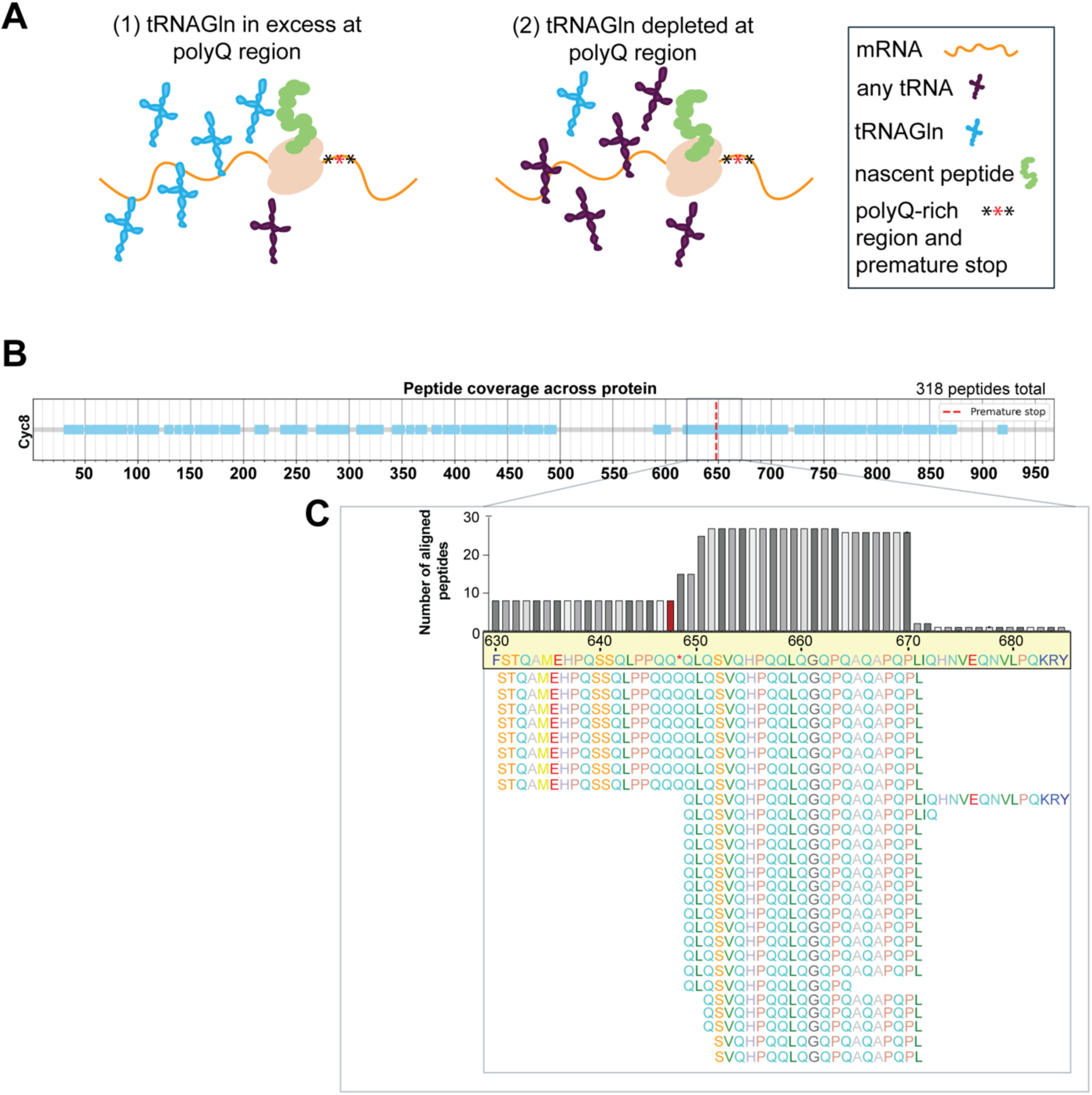
Glutamine is incorporated at the site of premature stop codon readthrough in Q-rich sequences. (A) Models of tRNA-based readthrough mechanisms. (B) Coverage plot showing the 318 chymotrypsin-digested peptides from replicate 1, detected by mass spectrometry and aligned to Cyc8, premature stop indicated by dotted red line. (C) Zoomed window showing chymotrypsin-digested peptide alignment of amino acids 630-686 with the number of reads for each residue plotted above. Location of premature stop highlight in red in plot.

### Proteins with short C-terminal glutamine repeats are enriched for strong stop codon contexts

Given that tRNA^Gln^ can decode stop codons in Q-rich sequence contexts, we filtered the yeast proteome to ask whether yeast proteins that naturally terminate within or adjacent to Q-rich sequences show a bias toward inherently stronger stop codon signals. We first examined whether [*PSI*^+^] status or canonically terminating QXQ context had a detectable effect on ribosome occupancy past canonical stop codons, finding that transcripts under either condition, [*PSI+*] transcriptome-wide or [*PSI+*] and all 4 QXQ-ending natural stops, did not show elevated 3’UTR ribosome density 100nts after the stop codon in our datasets (Figure 6A; Figure S6). Notably, only 4 transcripts contained the QXQ motif, limiting further investigation into these motifs (Figure S7). Furthermore, examination of the 264 yeast proteins ending in Q revealed that only 20 terminate in QQ or QQQ, and none displayed evidence of readthrough at their canonical stop compared to a random sample of 20 non-QQX ending proteins (Figure 7B).

**Figure 6.**
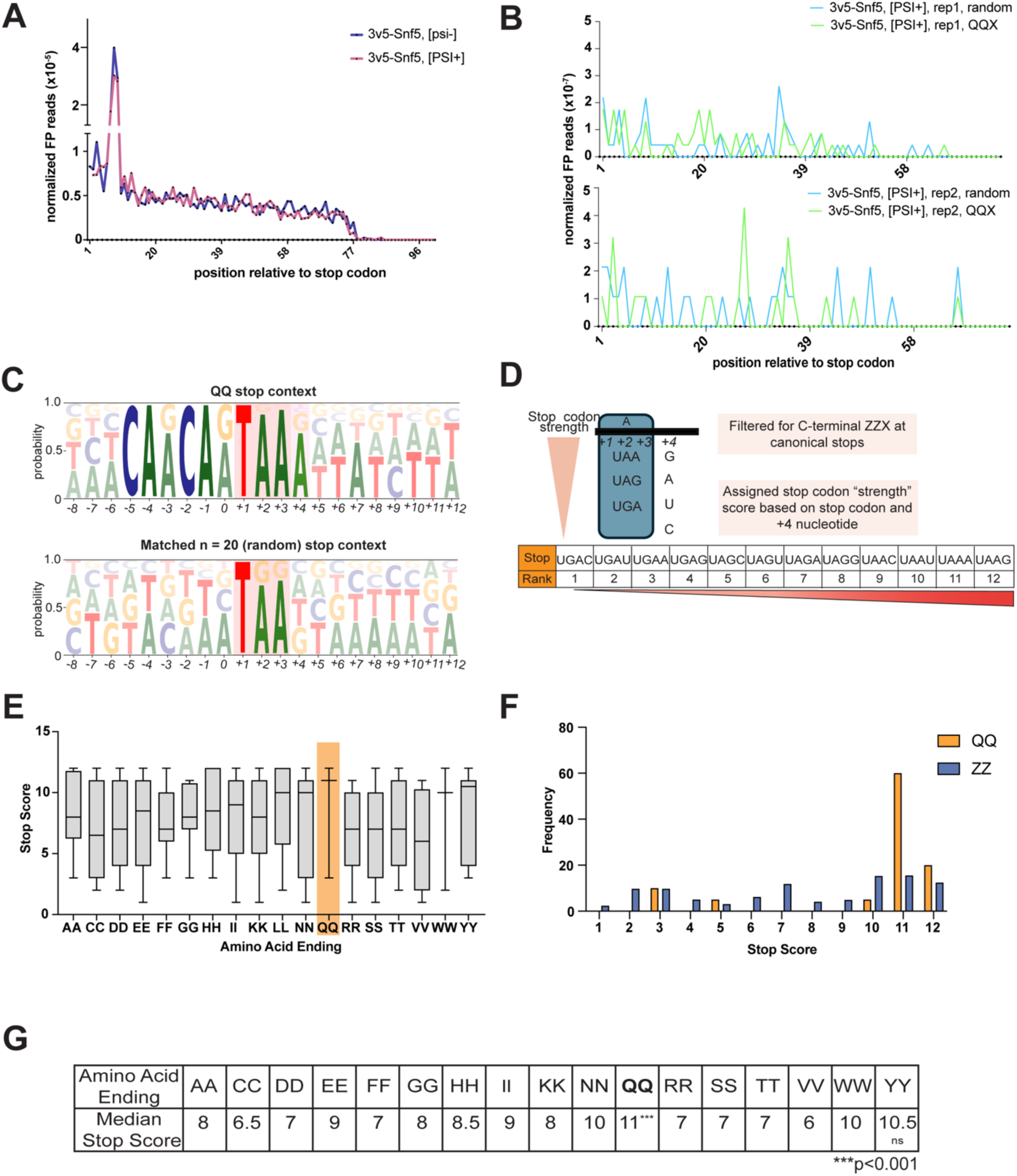
Proteins with short C-terminal glutamine repeats are enriched for strong stop codon contexts. (A) Quantification of global, normalized footprint reads 100nts after all canonical stop codons in [*PSI+*] and [*psi-*] *3v5-SNF5* samples. (B) Quantification of QQX (n=20) and a random sampling of n=20 transcripts (‘random’) normalized footprint reads after all canonical stop codons for [*PSI+*] samples only. Replicates shown in individual plots. (C) Natural stop codon context +/- 9 nucleotides for QQX (n = 20, top) and the matched random sample from 6B (n=20, bottom). (D) Schematic of strong stop signatures detailing filtering and ranking system for evaluating stop codon strength at QQX vs ZZX natural stops. (E) Box plot showing distribution of stop scores for all ZZX endings with n>3 proteins, QQX highlighted. (G) Histogram of stop scores of QQX proteins compared to all ZZX proteins. (G) Median stop scores for all ZZX endings with n>3 proteins. ‘QQ’ ending show significantly higher stop scores (Mann-Whitney U =7522.5, n_QQ = 20, n_ZZ = 515, p = 4.28 x 10^-4^, rank-biserial r=0.46). YYX endings were also tested but not significantly higher than all other (Mann-Whitney U = 2411.5, n_YY = 8, n_ZZ = 527, p = 0.482, rank-biserial = 0.14).

Closer examination of these QQ-terminating proteins revealed strong enrichment for TAA+4A stop codon context (Figure 7C), one of the most potent termination signals known. Quantitative ranking of stop+4 sequences using a previously published dataset ( (7); Figure 7D) confirmed that C-terminally QQ-containing proteins have significantly higher stop strength scores than all other duplicate C-terminal endings (Figures 7E–G). This evolutionary enrichment for strong termination signals at C-termini with short glutamine repeats is consistent with our finding that Q-rich sequences are inherently prone to readthrough and suggests that selective pressure has acted to reinforce stop codon fidelity at these loci.

## DISCUSSION

This study uncovers an unexpected connection between glutamine codon composition and premature stop codon readthrough. Our results demonstrate that a local QXQ motif broadly impairs termination fidelity, and that embedding this motif within a Q-rich sequence further promotes readthrough through tRNA^Gln^ insertion at near-cognate stop codons. The evolutionary bias for strong stop codon contexts at naturally glutamine-enriched protein termini reinforces the biological relevance of this finding. Together, these results define a codon-context-dependent mechanism of stop codon reinterpretation with broader implications for translational fidelity, and the design of therapeutic or synthetic constructs targeting premature stop codons.

### The QXQ motif broadly impairs termination fidelity

Our results complement prior findings by demonstrating that, across a range of QXQ variants tested outside of glutamine-rich domains, CAA flanking codons are more permissive for readthrough than CAG, consistent with the stronger near-cognate pairing potential of CAA at UAA and UAG stop codons. Although we did not determine the identity of the amino acid incorporated in the *ADE2* QXQ-only context, a prior study examining a QXQ-containing premature stop in the yeast protein Ste6 (which is not glutamine-rich, at ∼4% Q) found that tyrosine, lysine, and tryptophan were the predominant substitutions at the stop codon (19). Analysis of QXQ readthrough in human cells similarly revealed that, at this motif, tyrosine and glutamine are incorporated at roughly equal frequencies (20). Taken together with our data, these findings suggest that the QXQ motif broadly destabilizes termination, permitting competition by multiple near-cognate tRNAs at the stop codon rather than favoring insertion of a specific amino acid.

### Regional glutamine frequency drives tRNA^Gln^ misincorporation at the stop codon

The identity of the tRNA that miscodes at QXQ motifs may depend on the broader sequence environment. Our data show that readthrough at a QXQ motif is further elevated when the surrounding mRNA is enriched for glutamine codons. Mass spectrometry analysis of readthrough peptides from a glutamine-rich premature stop transcript detected glutamine insertion exclusively at the stop codon position, with no miscoding at flanking sites. This specificity suggests that regional codon composition influences which near-cognate tRNA successfully competes at the stop. Together with prior evidence that global tRNA availability modulates stop codon suppression (49–52), these data support a model in which active translation of glutamine-rich sequences locally enriches tRNA^Gln^, enabling it to outcompete both eRF1 and other near-cognate tRNAs at the stop codon.

### Eukaryotic ribosomes tolerate first-position mispairing at stop codons without downstream fidelity loss

The bacterial and eukaryotic decoding centers share extensive conformational similarities (53). In bacteria, miscoding at the P-site can propagate to the neighboring A-site, compromising decoding fidelity and triggering abortive termination by RF2 (47, 54). However, evidence from *in vitro* reconstituted yeast translational machinery suggests that a downstream loss of fidelity does not occur after a miscoding event in yeast (55). Our *in vivo* data further support this distinction between eukaryotic and prokaryotic translational systems by showing that readthrough in a glutamine-rich eukaryotic context involves misincorporation at the stop codon without detectable loss of fidelity at downstream positions. Our findings suggest that the eukaryotic translation machinery is tolerant of first-position mispairing events at stop codons, at least in the context of tRNA^Gln^ decoding, and that such errors do not propagate into the downstream coding sequence.

### Mechanistic basis of QXQ-mediated readthrough

How glutamine codons flanking the stop codon promote readthrough mechanistically remains an open question. None of the candidate quality control pathways we tested fully accounts for QXQ readthrough or the regional specificity we observe. Interestingly, however, deletion of NMD components increased readthrough without affecting transcript levels, suggesting a noncanonical translational role for the NMD machinery at premature stop codons that is supported by previous work (56).

One possibility towards the mechanism of QXQ readthrough is that tRNA^Gln^ isoacceptors possess structural or biochemical properties that favor engagement with the ribosomal decoding center, facilitating mispairing when tRNA^Gln^ occupies adjacent ribosomal sites. Structural data show that, in addition to eS30/Rps30b, the ribosomal protein uS19 contacts both A- and P-site tRNAs during decoding and may contribute to tRNA selection fidelity (53). Although we found no evidence for a role of the previously described eS30-tRNA^Gln^ interaction at the site we examined, it remains possible that eS30 or uS19 engages tRNA^Gln^ through alternative contacts in the decoding center. The translational initiation factor eIF3 has also been implicated in programmed stop codon readthrough via third-position miscoding (57, 58), though this is mechanistically distinct from the first-position mispairing we observe here. More broadly, resolving the mechanism underlying QXQ-mediated readthrough may ultimately shed light on why stop-to-glutamine codon reassignment has recurred independently across multiple eukaryotic lineages.

### Stop codon selection at Q-enriched termini reflects selective pressure against readthrough

Proteins with C-terminal extensions are generally deleterious, as such extensions can compromise protein stability or promote aggregation (59, 60). Transcripts are therefore expected to be under selective pressure to minimize readthrough, particularly when highly expressed or when their sequence context is inherently readthrough-permissive. Consistent with this, studies in bacteria, yeast, and humans have shown that highly expressed genes preferentially use UAA stop codons, suggesting that stop codon identity is subject to selection in contexts where readthrough is more likely (61–63). Our finding that proteins naturally terminating in QQ or QQQ motifs are strongly enriched for the TAA+4A stop codon context extends this principle to a specific sequence class. These data demonstrate that examining endogenous stop codons in the context of known readthrough-promoting motifs can reveal previously unappreciated patterns of stop codon selection shaped by the threat of translational readthrough.

### Implications for premature stop codon therapeutics

Premature stop mutations account for approximately 10-15% of all disease-causing genetic variants (30). QXQ motifs have been shown to elevate readthrough in diverse organisms including mammals (20, 21), suggesting that the sequence-context dependence of readthrough we describe in yeast is likely to extend to human disease contexts. Furthermore, engineered suppressor tRNAs have recently been shown to increase readthrough and alleviate disease phenotypes for a range of pathogenic premature stop mutations (64, 31, 32). Our results indicate that intrinsic readthrough propensity varies substantially depending on the local and regional sequence environment surrounding the premature stop, which may have significant implications for predicting the efficacy of both small-molecule readthrough compounds and suppressor tRNA-based therapeutic approaches.

## MATERIALS AND METHODS

### Yeast strains

All yeast strains were derived from the W303 background unless otherwise specified. Strains and genotypes are listed in Supplementary Table 1. W303 [*PSI+*] and [*psi-*] strains were kindly provided by Dan Jarosz (Stanford; UB33534 and UB33535 in Supplementary Table 1). Gene deletions were generated via Pringle-style insertion of a drug resistance cassette replacing the endogenous CDS (65). The premature stop alleles and wildtype controls were constructed in the following manner. First, the gene of interest, 500bp upstream and 500bp downstream were amplified from W303 genomic DNA. Second, the gene of interest was cloned via Gibson assembly (66) into a *LEU2* single integration vector with an N-terminal 3V5 tag, endogenous promoter (500bp upstream of gene) and endogenous 3’UTR (500bp downstream of gene). Third, the premature stop allele was generated via Q5 site-directed mutagenesis of the assembled plasmid, with the exception of *snf5-Q225X* and *snf5-Q267X*, which were amplified from the screen hits [(33)]. *CYC8* and *cyc8-Q648X* contain a missing 87nt from 1755nt-1842nt, deviating from W303 *CYC8*, due to a repetitive sequence that could not be PCR amplified.

The *RPS30B*/*RPS30-R10A*; *3V5-SNF5*/3V5-snf5-Q267X [*psi-*] and [*PSI+*] strains were generated by crossing [*psi-*] 74D-694 strains containing *rps30AΔ::natNT2 rps30B*Δ and YEplac112-*RPS30B-FLAG-TRP1* or YEplac112-*RPS30B-R10a-FLAG-TRP1* kindly provided by Leos Shivaya Valásek (Czech Academy of Sciences) into our W303 [*PSI+*] strains containing either *leu2::3V5-SNF5::LEU2* or *leu2::3V5-snf5-Q225X::LEU2*. These strains contained the same auxotrophies with variable mutant alleles, see Supplementary Table 1.

The *pTetO7.1-cyc8Q648X-3V5* plasmid was constructed via 3-piece Gibson Assembly with TetO7.1 from the WTC 846 system and the *cyc8-Q648X* premature stop allele into a LEU2 single integration vector, which was expressed with the partner *pRNR2* repressor plasmid in a *URA3* single integration vector ((48) ; *pRNR2*: Addgene plasmid #127575).

For all Gibson Assembly reactions, the NEBuilder HiFi Master Mix was used according to kit instructions (E5520S, *New England Biolabs*). All single integration plasmids were digested with PmeI or AscI before transformations and integrations were verified by PCR. All strains and plasmids used in this study are available upon request.

### Growth conditions

Unless otherwise stated, cells were grown overnight at 30°C in liquid YPD [1% yeast extract, 2% peptone, 2% dextrose, tryptophan (96 mg/L), uracil (24 mg/L), adenine (12 mg/L)] to saturation. The next day, cell density was quantified using a spectrophotometer, and cells were back-diluted to the indicated OD_600_ on the day of collection.

For profiling readthrough via immunoblot, cells were grown overnight in standard YPD conditions and back-diluted to an OD_600_ = 0.1 on the day of collection, then grown at 30°C until they reached OD_600_ ∼0.4-0.6, when immunoblotting samples were collected (see Immunoblotting).

For visualizing [*PSI+*] aggregates with NM-GFP, cells were grown overnight at 30°C in liquid synthetic complete media without supplemental uracil (0.67% yeast nitrogen base, 2% dextrose, custom dropout mix of adenine (20 mg/L), arginine (20 mg/L), methionine (20 mg/L), lysine (30 mg/L), tyrosine (30 mg/L), phenylalanine (50 mg/L), threonine (200 mg/L), tryptophan (20 mg/L), and histidine (20 mg/L)) to saturation. The day of imaging, cells were back-diluted to an OD_600_ = 0.2 in liquid synthetic complete media with raffinose and without supplemental uracil (0.67% yeast nitrogen base, 2% raffinose, custom dropout mix as listed above). Once cells reached OD_600_ ∼0.4-0.6, cells were pelleted and resuspended in liquid synthetic complete media with galactose and without supplemental uracil (0.67% yeast nitrogen base, 2% galactose, custom dropout mix as listed above). After 1 hour of galactose induction, cells were imaged (see Imaging NM-GFP aggregates).

For assaying turnover rate of 3v5-*ade2-E64QQ* and 3v5-*ade2-QE64QQ* using MG132 (Z-Leu-Leu-Leu-al, Sigma), cells were grown overnight in standard YPD conditions and back diluted to an OD_600_ = 0.2 on the day of collection. After cells reached OD_600_ = 0.5, the first (T_0_) timepoint was collected and then MG132 or DMSO was added to a final concentration of 100 µM for the remainder of timepoint collections.

For immunoprecipitation and mass spectrometry of cyc8-*Q648X* readthrough product, cells were grown overnight in standard YPD conditions. The day of collection, cells were back-diluted to an OD_600_ = 0.2 and once cells reached OD_600_ = 0.4, TetO-driven *cyc8-Q648X*-3V5 was induced by adding 2500 ng/mL of anhydrotetracycline (aTc, Cayman Chemical Company #10009542). 50 ODs of cells were collected when growth reached ∼OD_600_ = 1.0 and after ∼2-2.5 hours of aTc induction.

### Immunoblotting

Preparing lysates for SDS-Page was carried out according to Morse et al., 2024 (33). Briefly, 3 OD_600_ units were collected, resuspended in 5% trichloroacetic acid and incubated at 4 °C for at least 10 min to precipitate protein. Pellets were washed 1X with TE buffer and acetone, dried and lysed by bead beating in lysis buffer containing protease inhibitors. SDS buffer containing β-mercaptoethanol was added and samples were boiled for 5 min at 95 °C.

5 μl of samples and 3 μl of Precision Plus Protein Dual Color Standard (1610374, *Bio-Rad*) were loaded into 4–12% Bis-Tris Bolt gels (*Thermo Fisher Scientific*) and run at 150 V for 45 min. Protein was then transferred to 0.45 µm nitrocellulose membranes (*Bio-Rad*) with cold 1X Trans-Blot Turbo buffer in a semi-dry transfer apparatus (Trans-Blot Turbo Transfer System, *Bio-Rad*), or, if protein was over 100 kDa (Cyc8, Snf5), in a wet transfer apparatus for 3 h at 70V. Membranes were incubated at room temperature for 1 h in PBS Odyssey Blocking Buffer (927-4000, *LI-COR*) and incubated in primary antibody solutions at 4°C overnight. Membranes were then washed three times in 1X PBS with 0.1% Tween-20 (PBS-T, 5 min per wash) at room temperature before incubating in secondary antibody solutions at room temperature for 2.5 h. Membranes were washed three times in PBS-T at room temperature prior to imaging with the Odyssey system (*LI-COR Biosciences*).

All primary and secondary antibodies were diluted in PBS Odyssey Buffer with 0.1% Tween-20. Primary antibodies: mouse α-V5 antibody (R960-25, *Thermo Fisher Scientific*) used at a 1:2,000 dilution, rabbit α-hexokinase (100-4159, Rockland) used at 1:20,000 dilution. Secondary antibodies: goat α-mouse secondary antibody conjugated to IRDye 800CW used at a 1:15,000 dilution (926-32213, *LI-COR*); α-rabbit secondary conjugated to IRDye 680RD at a 1:15,000 dilution (926-68071, *LI-COR*).

### Growth curves

Growth curve data was collected as described in Morse *et al.* (33). Briefly, cells were back-diluted from a saturated overnight culture to an OD_600_=0.1 in liquid YPD in a 96-well plate in triplicate. Cells were grown at 30 °C for 24 h in a plate reader, and absorbance readings were collected every 15 min at A = 600nm.

### RNA extraction for RT-qPCR

RNA extraction was performed according to Morse et al., 2024 (33). Briefly, 2-4 mL of cells were collected by centrifugation and snap frozen in liquid nitrogen. Cells were thawed and resuspended, and RNA was phenol-chloroform extracted. After overnight – 20 °C precipitation in isopropanol and sodium acetate, RNA pellets were washed with 80% ethanol and resuspended in DEPC water. Total RNA was quantified by Nanodrop.

### Reverse transcription and qPCR

Reverse transcription and qPCR were carried out as described in Morse et al. (33). Briefly, 2.5 µg of RNA was treated with DNase using the TURBO DNA-free Kit (AM1907, *Invitrogen*). cDNA was synthesized from DNase-treated RNA via random hexamers and reverse transcription using the SuperScript^TM^ III (18080044, *Invitrogen*) according to manufacturer’s instructions. Quantitative PCR was performed using 5.2 μl Absolute Blue qPCR Mix (AB4162B, *ThermoFisher Scientific*), 100 nM of forward and reverse primer, and 4.8 μl of cDNA diluted 1:20 in nuclease-free water. Samples were run on a StepOnePlus (*Applied Biosystems*) qPCR machine in triplicate. Primers used for qPCR are listed in Supplementary Table 2.

### Imaging NM-GFP aggregates

3 µL of culture was placed on a slide and NM-GFP signal was imaged on a DeltaVision Elite widefield fluorescence microscope (GE Healthcare) running the accompanying softWoRx software (version 6.5.2), using a 60x/1.42 NA oil-immersion objective and a PCO Edge sCMOS camera. Z-stacks were acquired over 8 µm at 0.5 µm intervals (16 slices) using the FITC channel at 10% transmission with a 0.025 s exposure. A brightfield reference image was collected at each Z-slice with 32% transmission and 0.1 s exposure. Images were maximum projected in ImageJ and representative projections are shown.

### Frogging strains

Cells were grown overnight in standard YPD conditions and back-diluted to OD_600_ = 0.1 the next day. Once cells reached OD_600_ = 0.4-0.6, dilutions were performed. Cells were diluted to 0.1 OD/mL in 250 µL and diluted 5 more times in a 1:5 dilution series (6 total dilutions including 0.1OD/mL). 3 µL of each dilution was plated on synthetic complete plates without supplemental adenine (2% agar, 0.67% yeast nitrogen base, 2% dextrose, custom dropout mix of uracil, arginine, methionine, tryptophan and histidine at 20 mg/L, lysine and tyrosine at 30 mg/L, phenylalanine (50 mg/L), and threonine (200 mg/L)). Plates were imaged after 2 days of growth at 30 °C.

### Ribosome profiling and sequencing

Ribosome profiling collection and library preparation was performed as previously described for vegetative samples in *Powers, et al*. (67, 68). Briefly, cells were treated with cycloheximide for 30 s, filtered and flash frozen. During collection, an additional aliquot of cells was flash frozen to confirm each premature stop mutation via long read DNA sequencing of amplified PCR product, prior to library preparation. Extracts were milled under cryogenic temperatures and stored at –80 °C in aliquots. RNA was extracted from monosomes and ∼28-32 nt-size fragments were gel extracted. Libraries were prepped via linker ligation and rRNA fragments were depleted from samples using biotinylated anti rRNA oligos. The matched mRNA-seq libraries were prepared in parallel. These samples were Poly A-selected, fragmented and size selected for fragments ∼35-80 nts. No rRNA depletion was performed on mRNA-seq samples.

For slow growing 3V5-*snf5-Q225X* [*psi-*] strains, cells were back diluted in 5, 50mL cultures to OD_600_ = 0.2, collected when cultures reached OD_600_ ∼ 0.3-0.4 and pooled together once each culture’s *snf5-Q225X* mutation was verified. As less cells were collected for 3V5-*snf5-Q225X* [*psi-*] strains, half the amount (1.25 mL) of polysome lysis buffer was added prior to milling/cell lysis.

Samples were sequenced using 100 nt reads on a NovaSeq X Plus 10B Flow Cell at UCSF CAT, supported by UCSF PBBR, RRP IMIA, and NIH 1S10OD028511-01 grants. Barcoding primers used for library amplification are listed in Supplementary Table 2.

### Protein sample preparation for denaturing immunoprecipitation (IP)

50 OD of cells per replicate were pelleted in aliquots of 10 mLs. Pellets were resuspended in 4 mL 5% TCA and incubated at 4 °C overnight. Each 4 mL aliquot was then pelleted by centrifugation (20,000g x 1min), washed with 1mL acetone, then pelleted again (20,000g x 5min) and dried completely.

### Denaturing immunoprecipitation (IP) and mass spectrometry (MS)

Denaturing immunoprecipitation was performed as described in (Chen et al., 2020) with the following modifications. Briefly, dried pellets were resuspended in lysis buffer containing protease inhibitor and lysed on a bead beater using zirconia beads. 1% SDS was added and samples were boiled. Immunoprecipitation buffer was added to dilute SDS to 0.1%, cell material was pelleted and 97.5 µL anti-V5 agarose beads (Sigma) were added to 3,825 µL cleared lysate supernatant. Lysate:bead mixtures were incubated at 4 °C for 7 h with rotation. Beads were washed 2X with a high salt 1% NP-40 buffer and 4X with low-salt 0.05% NP-40 buffer. After the final washing step, the beads were resuspended in 1X SDS sample buffer (6.25 mM Tris, pH 6.8, 2% β-mercaptoethanol, 10% glycerol, 3% SDS, 0.0167% bromophenol blue) and boiled at 95C for 5 min to elute protein. 60 µL of sample and 3 μl of Precision Plus Protein Dual Color Standard (1610374, *Bio-Rad*) were loaded into separate lanes of a 10-well 4–12% Bis-Tris Bolt gel (*Thermo Fisher Scientific*) and run at 100 V for 90 min. *Cyc8-Q648X* readthrough product was visualized using a collodial blue straining kit (LC6035, Invitrogen); gel was stained for 3 h in colloidal blue solution and destained in milliQ water overnight. The next day, gel slices corresponding to Cyc8 readthrough product were extracted and submitted for chymotrypsin digestion and MS analysis at the Vincent J. Coates Proteomics/Mass Spectrometry Laboratory at the University of California, Berkeley.

In-gel chymotrypsin digestion was performed as follows: the gel slices were cut into 1 mm^2^ cubes with fresh razor blade and transferred to a clean microcentrifuge tube. The gel pieces were washed twice with 50% ACN and 50 mM Ammonium Bicarbonate pH 8 (ABC) for 15 min with shaking. The gel pieces were dehydrated with 100% ACN for 5 min with shaking. Then the solvent was removed, and the gel pieces were allowed to air dry for 5 min. Ten mM TCEP and 40 mM CAA were added to the dry gel pieces and incubated at 70 °C for 5 min. The gel pieces were washed again with 50% ACN and 50% 50mM ABC for 15 min with shaking, then dehydrated with 100% ACN for 5 min. After ACN, gel pieces were dried for 5 min. Gel pieces were rehydrated in 50 mM HEPES pH 8 (minimum volume, enough to cover pieces). Chymotrypsin (1ug) was added and after 1 hour of incubation at room temperature, 50mM HEPES pH 8 solution was added to cover the pieces and samples were incubated overnight at 37C. Peptides were extracted from gel pieces by washing twice with 25% ACN for 5 minutes with shaking, then once with 100% ACN for 5 minutes with shaking. Peptide extract solutions were filtered through a 0.22 µm PVDF spin column (Millipore). Peptide extract solutions were dried to 30-60 µL in a speedvac and acidified with 2 µl of formic acid (NEAT).

In-gel chymotrypsin digested peptides were analyzed by online capillary nano-MS/MS using a 25 cm reversed phase column and a 5mm precolumn (C18 300µmX5mmX5µm) (Thermo Fisher 174500). The separation column was fabricated in-house (75 µm inner diameter, packed with ReproSil-Gold C18-1.9 μm resin (Dr. Maisch GmbH)) that was equipped with a laser-pulled nanoelectrospray emitter tip. Peptides were eluted at a flow rate of 250 nL/min using a linear gradient of 2–50% buffer B in 140 min (buffer A: 0.05% formic acid and 5% acetonitrile in water; buffer B: 0.05% formic acid and 80% acetonitrile in water) in an Thermo Fisher Vanquish Neo nanoLC system operated in the trap-elute mode. Peptides were ionized using a FLEX ion source (Thermo Fisher) using electrospray ionization into a Fusion Lumos Tribrid Orbitrap Mass Spectrometer (Thermo Fisher Scientific). Data was acquired in orbi-orbi mode. Instrument method parameters were as follows: MS1 resolution, 120,000 at 200 m/z; scan range, 350−1600 m/z. MS2 resolution was 30,000. The top 20 most-abundant ions were subjected to collision-induced dissociation with a normalized collision energy of 35%, activation q 0.25, and precursor isolation width 2 m/z. Dynamic exclusion was enabled with a repeat count of 1, a repeat duration of 30 s, and an exclusion duration of 20 s. RAW files were analyzed using PEAKS (Bioinformatics Solution Inc) with the following parameters: semi-specific cleavage specificity at the C-terminal site of W, Y, F, L and M, allowing for partial cleavage (i.e. only one end of mappable peptides needed to show W, Y, F, L or M cleavage), precursor mass tolerance of 10 ppm, and fragment ion mass tolerance of 0.02 Daltons. Methionine oxidation and phosphorylation of STY was set as variable modifications and cysteine carbamidomethylation was set as a fixed modification. Peptide hits were filtered using a 1% FDR.

PEAKS alignment was then visualized in Geneious Prime (v2025.2.2). Figures were recreated from Geneious Prime alignment visualization. Peptide coverage was visualized using matplotlib (v3.7.5) in Python.

### Quantification and statistical analysis

Plotting and statistics for all bar charts, column plots, and scatterplots were performed using Prism Graphpad (v11.0.0) unless otherwise stated.

### Immunoblot quantification

Immunoblot quantification was performed by quantifying signals from bands in Image Studio (*LI-COR*). For all blots quantified in this study, raw V5 signal was normalized to raw hexokinase (Hxk2) signal (V5/Hxk2).

For readthrough quantification of premature stop alleles, each premature stop V5/Hxk2 ratio was normalized to the average of three wildtype V5/Hxk2.

For Ade2 time course experiments in Figure S2D, MG132-treated V5/Hxk2 values were normalized to DMSO-treated V5/Hxk2 at the appropriate timepoint (MG132/DMSO) to quantify protein stability in the absence of proteosome degradation, then normalized values for each timepoint were plotted.

For Cyc8 time course experiments in Figure S5A, normalized TetO-driven *cyc8-Q648X*-3v5 readthrough product for each timepoint was expressed as a ratio to normalized Cyc8 wildtype protein driven by its endogenous promoter (lanes 1 and 2).

### qPCR Quantification

The C_T_ mean from each primer pair directed against the experimental target was normalized using the ΔC_T_ method. Statistical tests were performed on ΔΔC_T_ values.

### mRNA-seq and ribosome profiling analysis

Adapter sequences were trimmed, reads were filtered to exclude rRNA and aligned to a STAR index of the S288c R641-1.110 genome assembly as previously described in (69). The premature stop readthrough analysis in Figure 4 used only ribosome profiling reads filtered to 28-29 nts, as this is the expected ribosome-protected fragment size for cycloheximide-treated ribosomes and these sized reads had the cleanest frame distribution in our dataset across reads 26-34 nts [Figure S4F; (70)]. Ribosome profiling reads were A-site corrected based on fragment length. For mRNA-seq data, reads per ORF were counted and each gene’s RPKM was calculated by normalizing the raw reads to the total sum of reads per sample and the gene length.

Ribosome profiling data of individual transcripts was extracted from the corrected A-site file based on gene coordinates and each read was normalized to that sample’s transcript mRNA-seq RPKM (plotted in Figure 4B-D; S4I-J). In figures S4G-H, frame periodicity was defined by position 0 (first), 1 (second) or 2 (third) from the start nucleotide and the premature stop codon was included in the ‘after stop’ partition.

Ribosome readthrough score (RRTS) was calculated for all transcripts based on previously reported methods (40). Briefly, for each pre-defined region, ribosome densities were summed, adjusted to transcript length and converted to reads per kilobase (RPK). The region from the 5’ start site and the premature stop was defined as the ‘CDS’ and the region from the premature stop to the canonical 3’ stop codon was defined as the ‘3’UTR’. Coordinates for start and premature stop sites were included in ‘CDS’ and coordinates for canonical stop site was included in ‘3’UTR’. RRTS was calculated for both the wildtype and premature stop transcript by dividing the 3’UTR value by the CDS value.

The metagene plot in Figure 6A was created by taking 28-29 nt global A-site corrected values +100nt after the stop and dividing them by the total mapped reads for frame sizes 28-29nts. The metagene plots in Figure 6B were created in a similar manner, except all frame sizes were used due to lower n, and the random sample of n=20 proteins is used for 6B and 6C.

### Filtering for C-terminal Q-enriched and QXQ ending ORFs

First, a whole genome summary of stop codon context (stop codon and +/- 9 nucleotides of stop codon) was created using the S288c R641-1.110 genome assembly. Next, the yeast proteome was downloaded from Uniprot and filtered for proteins that ended in single, duplicate and triplicate glutamines (proteome downloaded 09/20/2024). Stop codon context was compared between QQX proteins and a random sample of n=20 proteins using logomaker (v0.8) in Python.

To find QXQ stops, the single glutamine-ending proteins were filtered using the whole genome summary of stop codon context for those that contained ‘CAA’ or ‘CAG’ as the codon immediately following the stop codon. We used IGV (2.18.0) to visualize ribosome density in and downstream of the CDS from the corrected A-site file for transcripts with QXQ stops.

To match duplicate amino acid endings, the yeast proteome was filtered for all proteins ending in duplicate amino acids and their sequence context information was extracted from the whole genome summary of stop codon context. From there, the stop codon and +4 nucleotide of each transcript was assigned a score based on a previous study in yeast (7). Files for whole genome stop context and duplicate amino acid filtering are available upon request.

### Statistical comparison of stop codon context scores

To test whether the distribution of stop codon context scores differed between ZZ-ending and QQ-ending genes, a two-sided Mann-Whitney U test was performed (Scipy 1.10.1, Python 3.8.5). This non-parametric test was chosen because the score variable is discrete (ranging from 1 to 12) and not normally distributed. A significance threshold of a=0.05 was used. Effect size was estimated using the rank biserial correlation (r = 1-2U/n_1_n_2_).

## Resource availability

All reagents used in this study are available upon request from the corresponding author. Sequencing data generated in this study will be publicly deposited at NCBI GEO and text will be updated when accession number is acquired.

## Author contributions

J.L., conceptualization, data curation, formal analysis, investigation, methodology, validation, visualization, and drafting of the manuscript. K.M. conceptualization, data curation, investigation, methodology, and validation. S.S., data curation, formal analysis, investigation, methodology, and validation. G.B., methodology, visualization, project administration. E.Ü., conceptualization, methodology, visualization, supervision, project administration, funding acquisition, and drafting of the manuscript.

## Acknowledgements

We thank Nicholas Ingolia, Kathleen Collins, Cyrus Ruediger, Ben Styler, Tianyao Xiao, Alex Wooldredge, Alejandro Collins, Camila Sousa, Claudia Medrano and Silvan Spiri for suggestions and comments on this manuscript. A special thanks to Nicholas Ingolia for his thoughtful advice on experimental directions and ribosome profiling methodology. We also thank Daniel Jarosz and Leos Shivaya Valásek for kindly gifting us yeast strains for the study. Finally, thanks to Kathleen Collins, Nicholas Ingolia, Daniel Jarosz, Jeremy Thorner and all members of the Brar and Ünal labs for valuable discussions. This work is supported by funds from the National Institutes of Health (R01 GM140005) and Astera Institute to EÜ.

