## Supplemental Materials for "Glutamine codon-driven translational readthrough reveals context-dependent stop codon decoding fidelity"

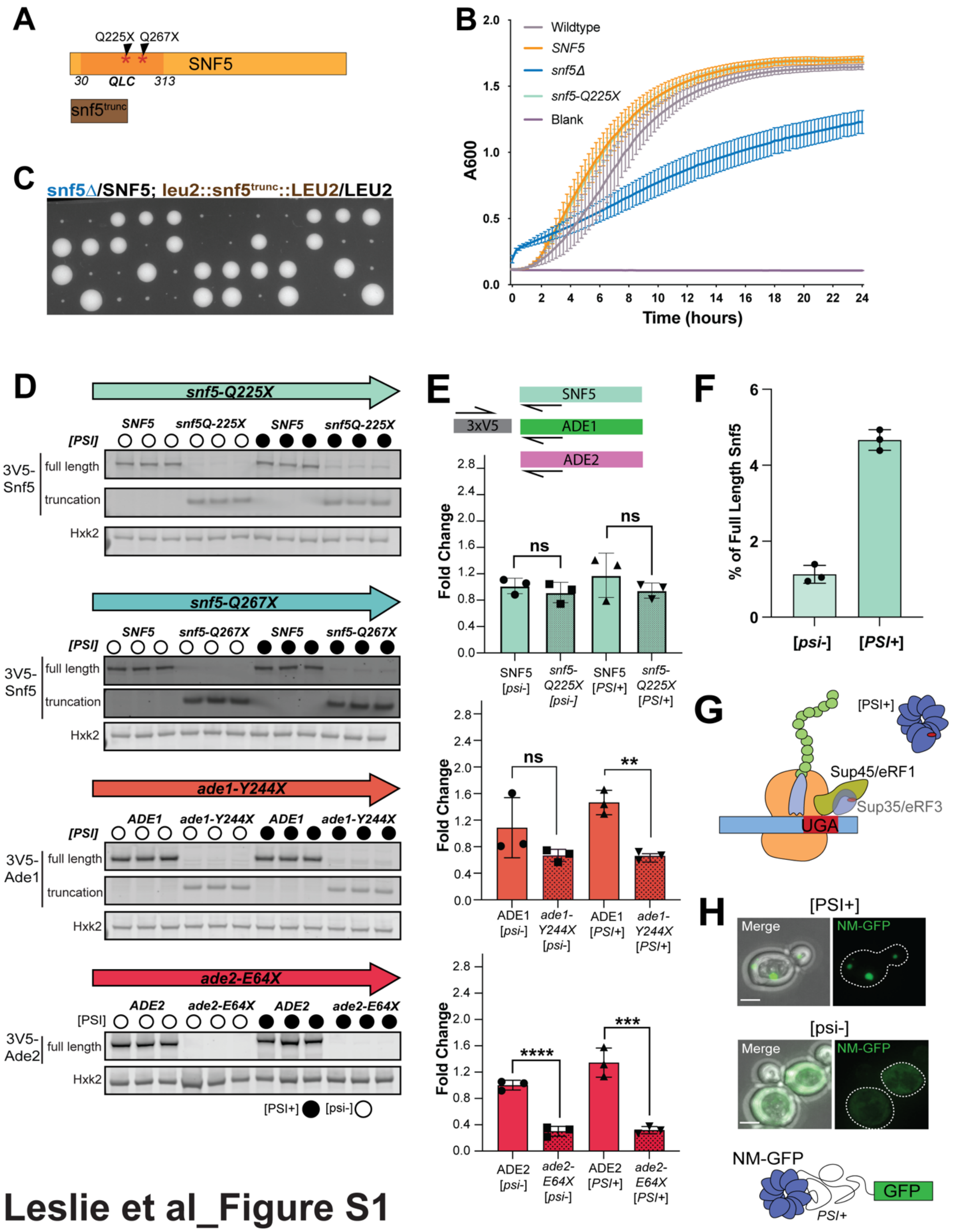


Leslie et al_Figure S1

**Figure 1 – Supplementary Data.** (A) Schematic of Snf5 premature stop alleles (top) and visual representation of *snf5*$^{trunc}$ (below). (B) Growth curves for cells grown in rich media with 2% sucrose. Absorbance readings at 600nm collected at 15-minute intervals for 24 h is plotted for wildtype (gray), SNF5-Tg (orange), *snf5*∆ (blue), *snf5-Q225X*-Tg (cyan). (C) Tetrad dissection of hemizygous *snf5*$^{trunc}$ strain. (D) Second technical replicate of semi-quantitative immunoblot replicates of 3v5-*snf5Q225X*, 3v5-*snf5Q267X*, 3v5-*ade1Y244X* and 3v5-*ade2E64X*. Average of technical replicate 1 (1C) and 2 plotted in 1E. (E) Fold change of *3v5-SNF5/3v5-snf5-Q225X*, *3v5-ADE1/3v5-ade1-Y244X* and *3v5-ADE2/3v5-ade2-E64X* transcripts in [*PSI*+]/[*psi*-] cells, as measured by RT-qPCR (primers as indicated). Statistical significance was determined by a two-tailed Welch's t-test on $\Delta\Delta$ $C_T$ values (**p<0.01, ***p<0.001 and ****p<0.0001). (F) Comparison of [*PSI+]* and [*psi*-] readthrough for *snf5-Q225X.* Average of two technical replicates from three biological replicates plotted. (G) Schematic of translational termination factors Sup45/eRF1 and Sup35/eRF3 bound to stop codon with [*PSI*+] aggregates that sequester soluble Sup35. (H) NM-GFP imaging to visualize presence or absence of [PSI+] aggregates for [*PSI*+] and [*psi*-] strains used in the study.

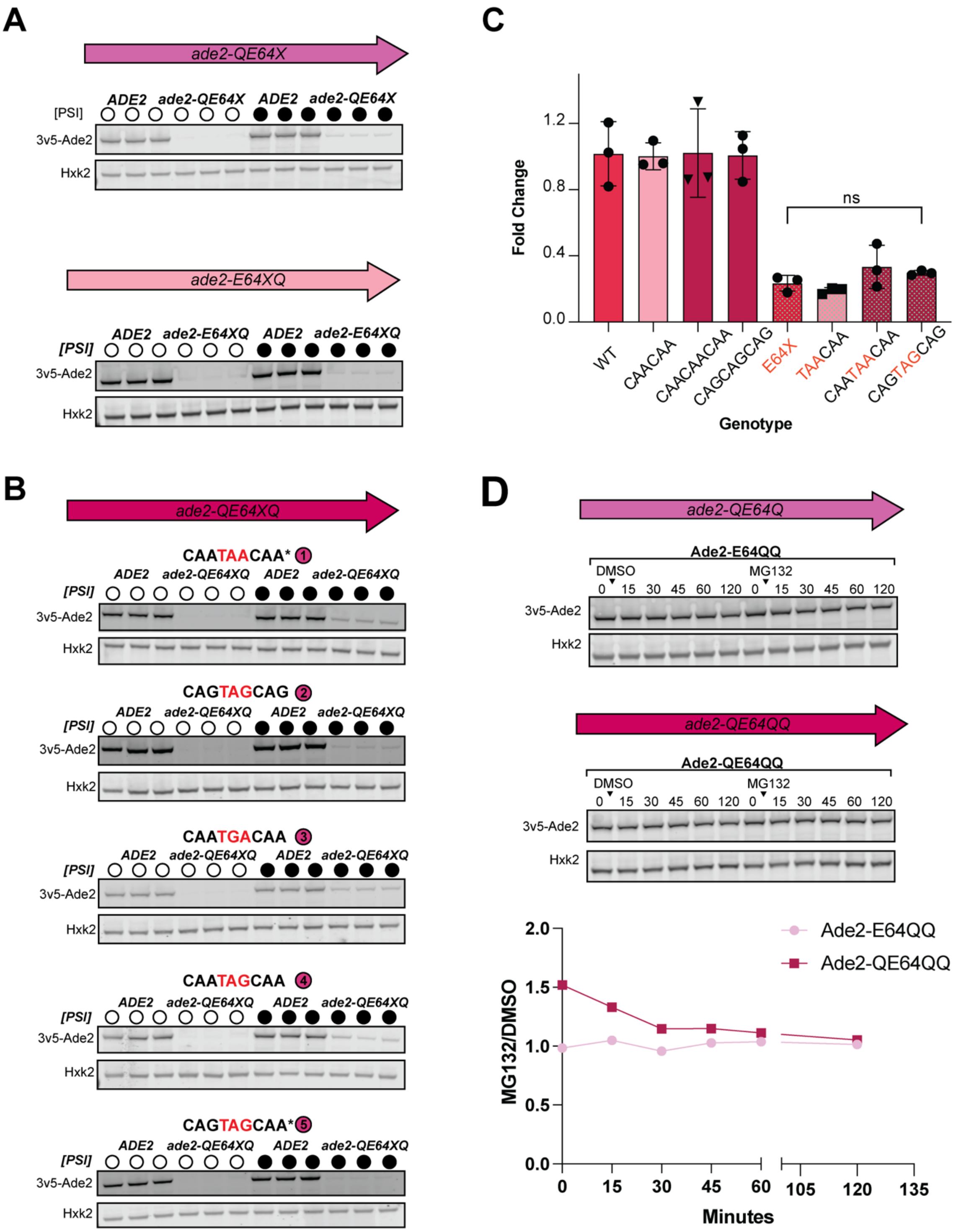


Leslie et al_Figure S2

**Figure 2 – Supplementary Data.** (A) Second technical replicate of immunoblots for Ade2 5' or 3' Q context premature stop alleles: CAA-TAA, TAA-CAA. Average of technical replicate one (2B) and two plotted in 2D. (B) Second technical replicate of immunoblots for Ade2 5' and 3' Q context premature stop alleles: CAA-TAA-CAA, CAG-TAG-CAG, CAA-TGA-CAA, CAA-TAG-CAA and CAG-TAG-CAA. Average of technical replicate one (2C) and two plotted in 2D. (C) Fold change of [*PSI*+] 3x-V5 tagged *ADE2* variant transcripts in [*PSI*+]/[*psi*-] cells profiled in S2A-B, as measured by RT-qPCR using primers indicated for ADE2 in S1E. All premature stop alleles are normalized to their respective WT allele set to 1. Error bars represent SD of n=3 biological replicates. No significant difference was observed among premature stop alleles by one-way ANOVA on $\Delta\Delta$Ct values of premature stop alleles, $F(3,8) = 2.991$, $p=0.0957$. (D) Representative semi-quantitative immunoblots showing time courses of Ade2 abundance after 100μM MG132 addition or DMSO control in *pdr5*Δ cells to measure turnover rate of Ade2 with Q substitutions as indicated. Quantification of MG132/DMSO Hxk2 normalized Ade2 abundance over time.

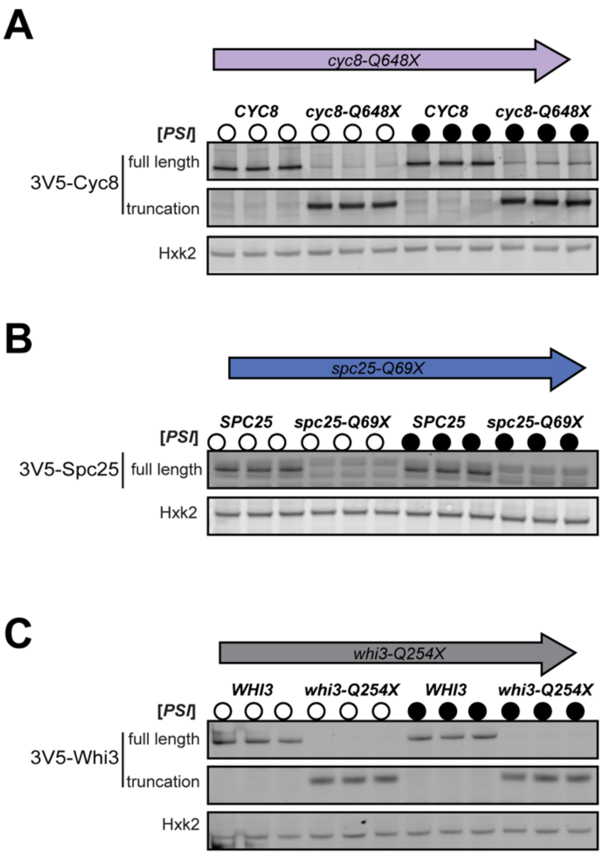


Leslie et al_Figure S3

**Figure 3 – Supplementary Data.** Second technical replicate of semi-quantitative immunoblot replicates of (A) *CYC8/cyc8Q648X* (B) *SPC25/spc25Q69X* (C) *WHI3/whi3Q254X*.

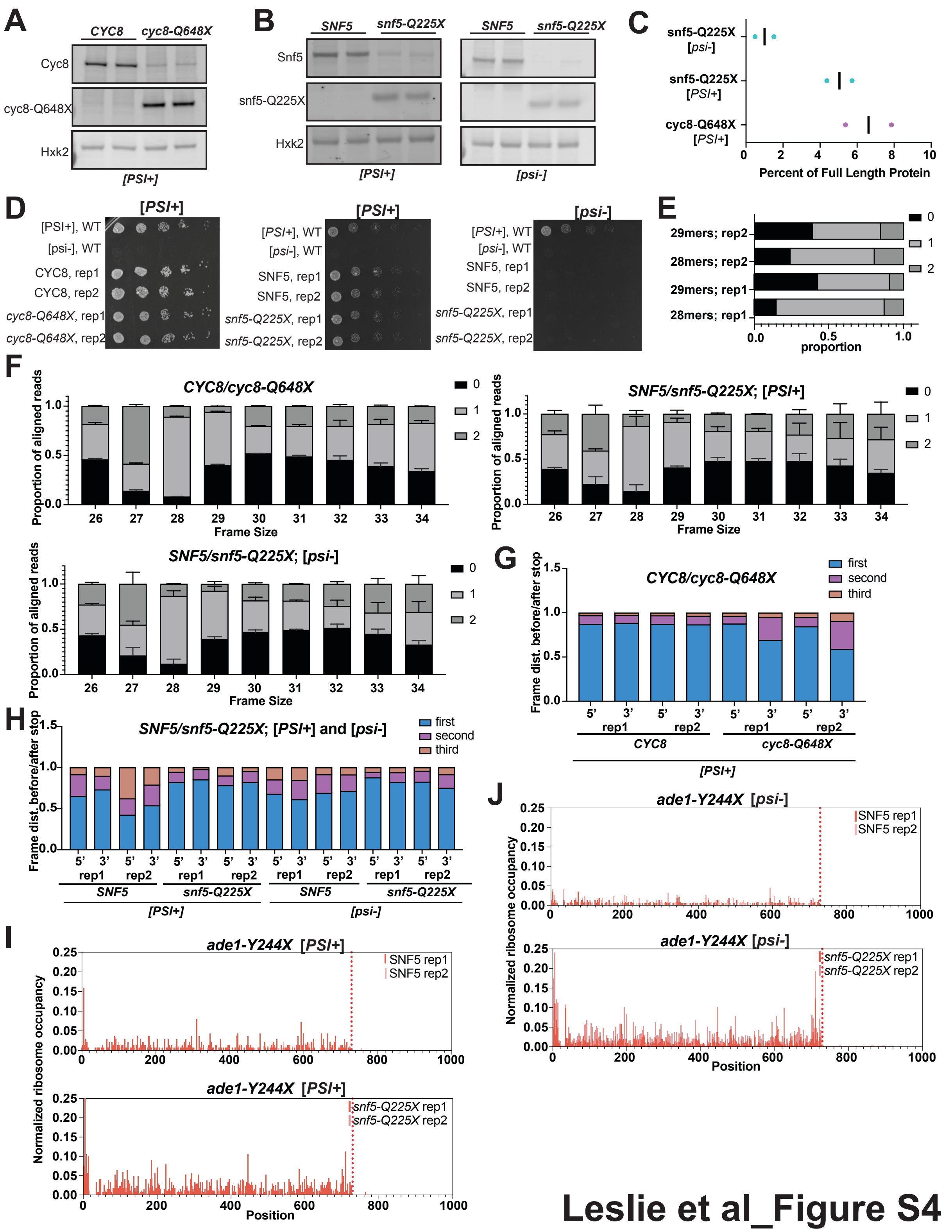


**Leslie et al_Figure S4**

**Figure 4 – Supplementary Data.** (A) Immunoblot of ribosome profiling [*PSI*+] strains 3v5-*CYC8*/3v5-*cyc8Q648X*. Protein collected from ribosome profiling cultures. (B) Immunoblot of ribosome profiling [*PSI*+] strains 3v5-*SNF5*/3v5-*snf5Q225X* (top) and ribosome profiling [*psi*-] strains 3v5-*SNF5*/3v5-*snf5Q225X* (bottom). Protein collected from ribosome profiling cultures. (C) Quantification of full-length protein produced from premature stop containing transcripts (A)-(C). (D) Ribosome profiling strains plated on synthetic complete media lacking adenine to confirm [*PSI*] status. (E) Distribution of aligned reads by p-site for Snf5 [PSI+] replicates. Rep2 shows lower bias towards a single frame, likely amounting to lower first frame distribution in (I). (F) Distribution of frames for each frame sizes 26-34nt. Cyc8/*cyc8Q648X* (left), Snf5/*snf5Q225X* [*PSI*+] (right) and Snf5/*snf5Q225X* [*psi*-] (bottom left). (G) Distribution of frame periodicity for 5' (before) and 3' (after) of premature stop CYC8/*cyc8Q648X.* (H) Distribution of frame periodicity for 5' (before) and 3' (after) of premature stop SNF5/*snf5Q225X* [*PSI*+] and SNF5/*snf5Q225X* [*psi*-]. (I) Normalized ribosome profiling traces of *SNF5/snf5*-*Q225X* [*PSI*+] *ade1-Y244X* controls. *ade1-Y244X* stop = 729nt. Two replicates plotted. (J) Normalized ribosome profiling traces of *SNF5/snf5*-*Q225X* [*psi*-] *ade1-Y244X* controls. *ade1-Y244X* stop = 729nt. Two replicates plotted.

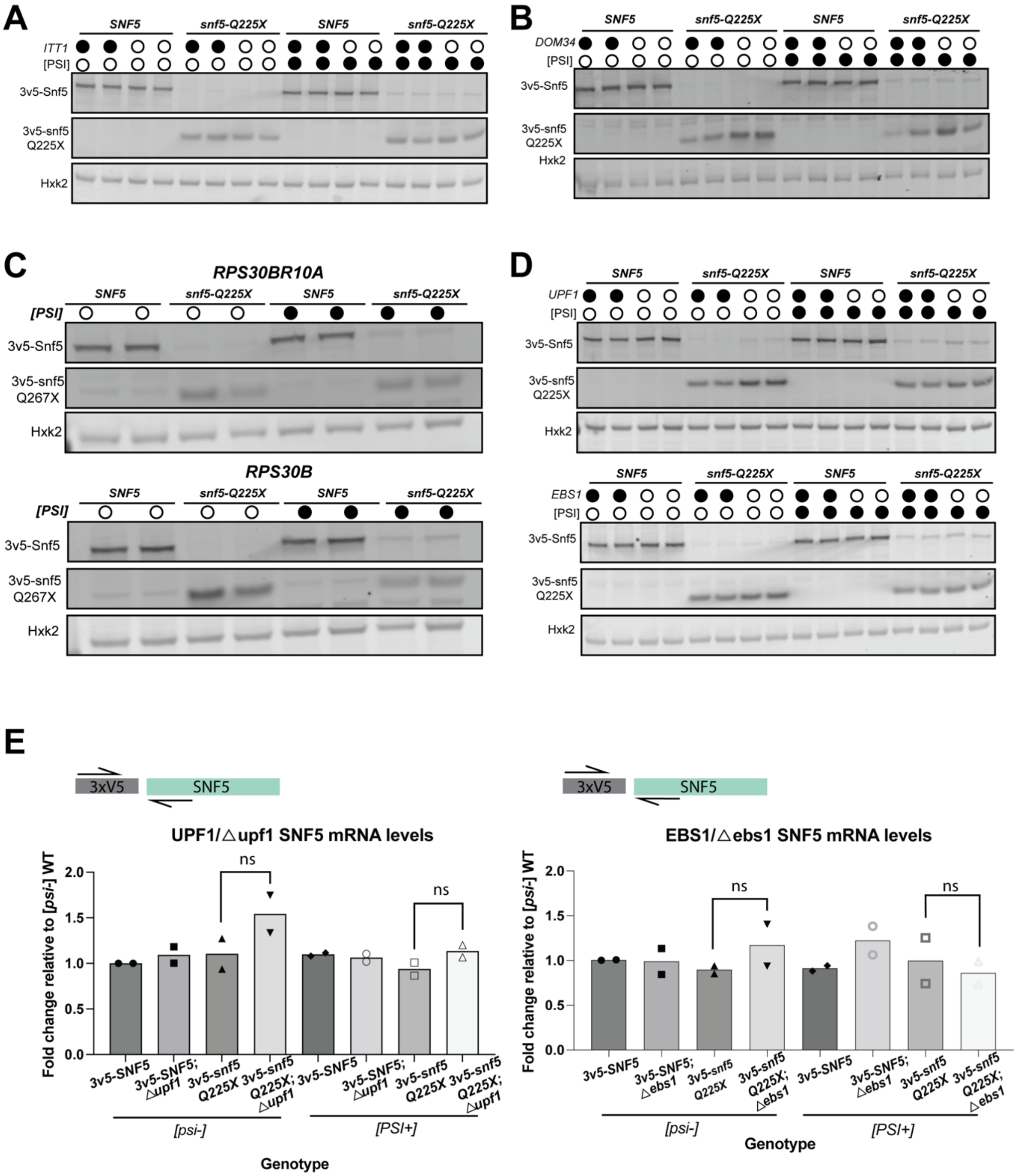


### Leslie et al_Supplementary Data Table 1

**Table 1– Supplementary Data.** (A) Representative immunoblot of *3v5-SNF5/3v5-snf5Q225X* readthrough in wildtype and *dom34*∆ mutants. [*PSI*+]/[*psi*-] conditions (as indicated). Quantified in Table 1. (B) Representative immunoblot of *3v5-SNF5/3v5-snf5Q225X* readthrough in wildtype and *itt1*∆ mutants. [*PSI*+]/[*psi*-] conditions (as indicated). Quantified in Table 1. (C) Representative immunoblot of *3v5-SNF5*/3v5-*snf5Q267X* readthrough in RPS30B-R10A mutant (top) and RPS30B control, [*PSI*+]/[*psi*-] conditions as indicated. (D) Representative immunoblot of 3v5-*SNF5*/3v5-*snf5Q225X* readthrough in wildtype and *upf1*∆ mutants, [*PSI*+]/[*psi*-] conditions as indicated. (E) Quantification (via qPCR) of [*psi*-] UPF1/*upf1*∆ *SNF5/snf5-Q225X* transcript abundance (left) and [*PSI*+] UPF1/*upf1*∆ *SNF5/snf5-Q225X* transcript abundance (right) in (A), followed by identical EBS1/ebs1∆ quantification. Statistical significance was determined using an unpaired two-tailed T-test on ∆∆Ct values, $^{*}p_{adj}<0.05$, $^{ns}p>0.05$.

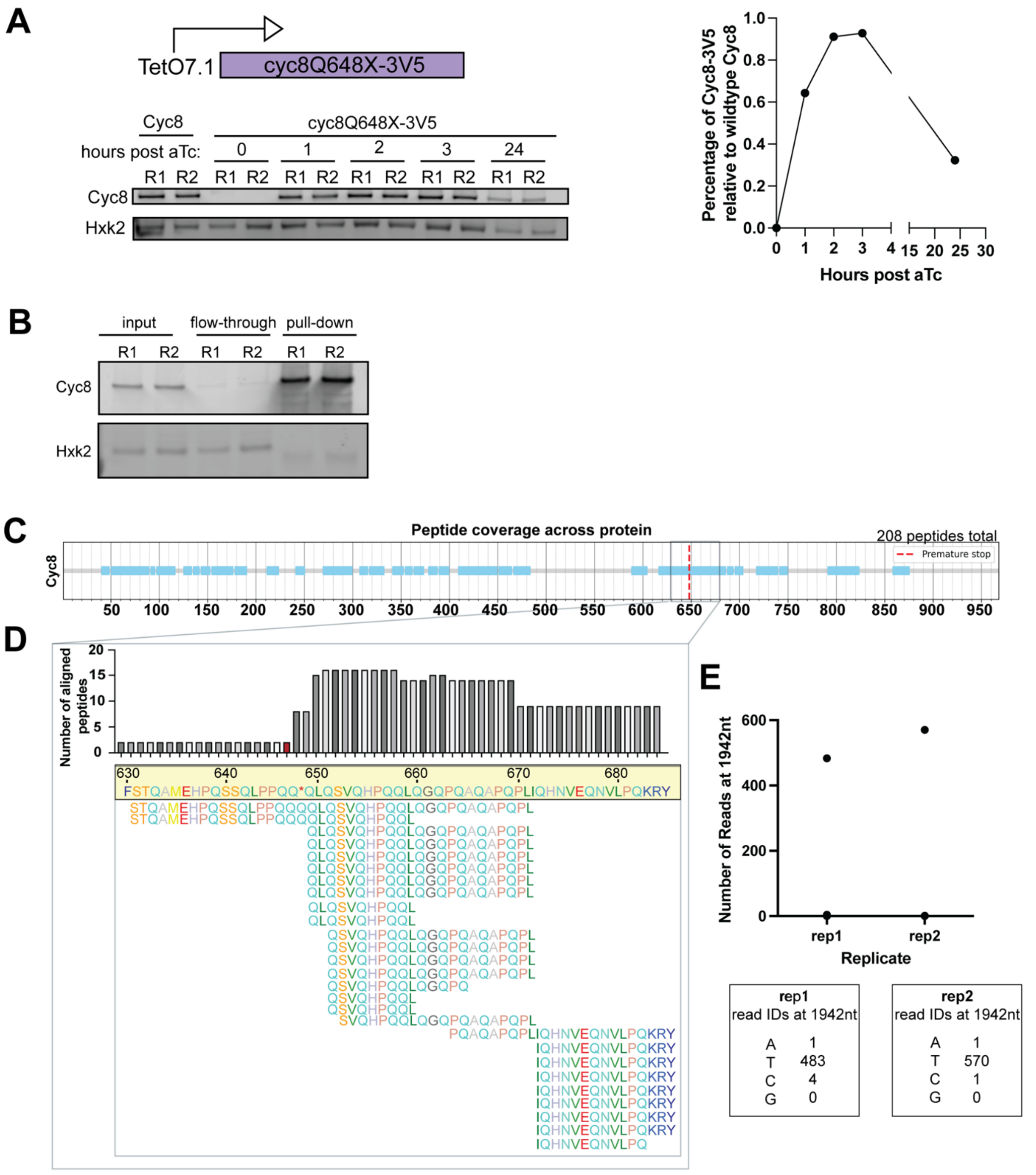


Leslie et al_Figure S5

**Figure 5 – Supplementary Data.** (A) Top - schematic of $P_{TetO7.1}$ controlled, anhydrotetracycline (aTc)-induced temporal overexpression of *cyc8Q648X*-3v5. Bottom - immunoblot timecourse data showing endogenous promoter Cyc8 expression levels compared to *cyc8-Q648X* post-induction for two replicates, hours post-induction as marked. Right - Quantification of $P_{TetO7.1}$ *cyc8Q648X* induction relative to wildtype Cyc8 expression over time. (B) Immunoblot of *cyc8-Q648X* immunoprecipitation samples (input, flow-through and final pulldown) submitted for mass spectrometry analysis. (C) Coverage plot showing the 208 chymotrypsin digested peptides from replicate 2, detected by mass spectrometry and aligned to Cyc8, premature stop indicated by dotted red line. (D) Zoomed window showing alignment of amino acids 630-686 with the number of reads for each residue plotted above. (E) Long read sequencing results from PCR amplified genomic DNA of mass spectrometry submissions. Read count plotted and read results for nucleotide 1942 (T of TAA premature stop) as indicated.

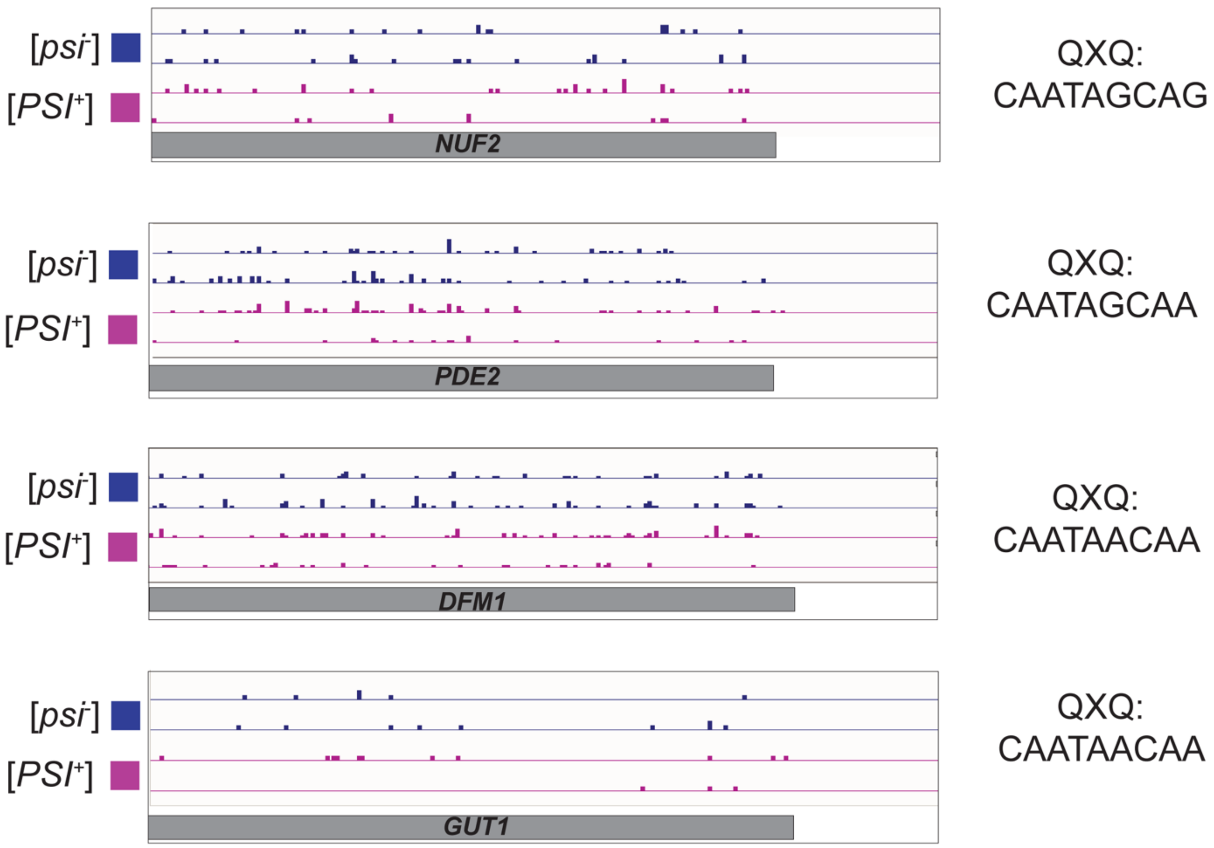


**Leslie et al_Figure S6**

**Figure 6 – Supplementary Data.** Ribosome profiling traces of 200nts upstream of QXQ natural stops (Nuf2, Pde2, Dfm1 and Gut1) and 50nts downstream from [*PSI*+] and [*psi*-] collections.

**Supplementary Table 1. Genotypes of the strains used in this study.**

| Strain | Genotype | Figure |
|---|---|---|
| UB18345 | MAT***a***, *ADE2, leu2-3, ura3, trp1-1, his3-11,15, can1-100, GAL, phi+*<br>MATα, *ADE2, leu2-3, ura3, trp1-1, his3-11,15, can1-100, GAL, phi+*<br>*snf5::KANMX* | 1A |
| UB18467 | MAT***a***, *ADE2, leu2-3, ura3, trp1-1, his3-11,15, can1-100, GAL, psi+*<br>MATα, *ADE2, leu2-3, ura3, trp1-1, his3-11,15, can1-100, GAL, psi+*<br>*snf5::KANMX leu2::SNF5::LEU2* | 1A |
| UB18469 | MAT***a***, *ADE2, leu2-3, ura3, trp1-1, his3-11,15, can1-100, GAL, psi+*<br>MATα, *ADE2, leu2-3, ura3, trp1-1, his3-11,15, can1-100, GAL, psi+*<br>*snf5::KANMX*<br>*leu2::snf5-Q225X::LEU2* | 1A |
| UB33534 | MATα, *leu2-3,-112; his3-11,-15; trp1-1; ura3-1; ade1-4; can1-100, PSI+, CAN1-370-TAA* | S4E-G |
| UB33535 | MATα, *leu2-3,-112; his3-11,-15; trp1-1; ura3-1; ade1-4; can1-100, psi-, pin-, CAN1-370-TAA* | S4E-G |
| UB37419 (MATα)<br>UB37420 (MATα)<br>UB37421 (MAT**a**) | *leu2-3,-112; his3-11,-15; trp1-1; ura3-1; ade1-4; can1-100, psi-, pin-, CAN1-370-TAA*<br>*leu2::3V5-SNF5::LEU2* | 1C; 1E; S1D; S1E |
| UB37422 (MAT**a**)<br>UB37423 (MATα)<br>UB37424 (MATα) | *leu2-3,-112; his3-11,-15; trp1-1; ura3-1; ade1-4; can1-100, psi-, pin-, CAN1-370-TAA*<br>*leu2::3V5-snf5-Q225X::LEU2* | 1C; 1E; S1D; S1E |
| UB37438 (MATα)<br>UB37439 (MATα)<br>UB37440 (MAT**a**) | *leu2-3,-112; his3-11,-15; trp1-1; ura3-1; ade1-4; can1-100, PSI+, CAN1-370-TAA*<br>*leu2::3V5-SNF5::LEU2* | 1C; 1E; S1D; S1E |
| UB37441 (MAT**a**)<br>UB37442 (MATα)<br>UB37443 (MATα) | *leu2-3,-112; his3-11,-15; trp1-1; ura3-1; ade1-4; can1-100, PSI+, CAN1-370-TAA*<br>*leu2::3V5-snf5-Q225X::LEU2* | 1C; 1E; S1D; S1E |
| UB39534 (MAT**a**)<br>UB39535 (MATα)<br>UB39536 (MAT**a**) | *leu2-3,-112; his3-11,-15; trp1-1; ura3-1; ade1-4; can1-100, psi-, pin-, CAN1-370-TAA*<br>*leu2::3v5-snf5-Q267X::LEU2* | 1C; 1E; S1D; S1E |
| UB39531 (MAT**a**)<br>UB39532 (MATα)<br>UB39533 (MATα) | *leu2-3,-112; his3-11,-15; trp1-1; ura3-1; ade1-4; can1-100, PSI+, CAN1-370-TAA*<br>*leu2::3v5-snf5-Q267X::LEU2* | 1C; 1E; S1D; S1E |
| UB37518 (MATα)<br>UB37519 (MAT**a**)<br>UB37520 (MATα) | *leu2-3,-112; his3-11,-15; trp1-1; ura3-1; ade1-4; can1-100, psi-, pin-, CAN1-370-TAA*<br>*leu2::3V5-ADE1::LEU2* | 1D; 1E; S1D; S1E |

| | | |
|---|---|---|
| UB37521 (MAT$\alpha$)<br>UB37522 (MAT**a**)<br>UB37523 (MAT**a**) | *leu2-3,-112; his3-11,-15; trp1-1; ura3-1; ade1-4; can1-100, psi-, pin-, CAN1-370-TAA leu2::3V5-ade1-Y244X::LEU2* | 1D; 1E; S1D; S1E |
| UB37447(MAT**a**)<br>UB37448 (MAT$\alpha$)<br>UB37449 (MAT$\alpha$) | *leu2-3,-112; his3-11,-15; trp1-1; ura3-1; ade1-4; can1-100, PSI+, CAN1-370-TAA leu2::3V5-ADE1::LEU2* | 1D; 1E; S1D; S1E |
| UB37444 (MAT**a**)<br>UB37445 (MAT$\alpha$)<br>UB37446 (MAT**a**) | *leu2-3,-112; his3-11,-15; trp1-1; ura3-1; ade1-4; can1-100, PSI+, CAN1-370-TAA leu2::3V5-ade1-Y244X::LEU2* | 1D; 1E; S1D; S1E |
| UB38470 (MAT$\alpha$)<br>UB38471 (MAT**a**)<br>UB38472 (MAT**a**) | *leu2-3,-112; his3-11,-15; trp1-1; ura3-1; ade1-4; can1-100, psi-, pin-, CAN1-370-TAA leu2:: 3V5-ADE2::LEU2* | 1D; 1E; S1D; S1E |
| UB38473 (MAT$\alpha$)<br>UB38474(MAT**a**),<br>UB38475 (MAT**a**) | *leu2-3,-112; his3-11,-15; trp1-1; ura3-1; ade1-4; can1-100, psi-, pin-, CAN1-370-TAA leu2::3V5-ade2-E64X::LEU2* | 1D; 1E; S1D; S1E |
| UB38476 (MAT$\alpha$)<br>UB38477 (MAT**a**)<br>UB38478 (MAT**a**) | *leu2-3,-112; his3-11,-15; trp1-1; ura3-1; ade1-4; can1-100, PSI+, CAN1-370-TAA leu2::3V5-ADE2::LEU2* | 1D; 1E; S1D; S1E |
| UB38479 (MAT**a**)<br>UB38480 (MAT**a**)<br>UB38481 (MAT$\alpha$) | *leu2-3,-112; his3-11,-15; trp1-1; ura3-1; ade1-4; can1-100, PSI+, CAN1-370-TAA leu2::3V5-ade2-E64X::LEU2* | 1D; 1E; S1D; S1E |
| UB28238 | MAT**a***, ADE2, leu2-3, ura3, trp1-1, his3-11,15, can1-100, GAL, phi+*<br>MAT$\alpha$*, ADE2, leu2-3, ura3, trp1-1, his3-11,15, ca1-100, GAL, phi+*<br>*snf5::KANMX leu2::snf5*$^{trunc}$*::LEU2* | S1C |
| UB27898 | MAT**a***, ADE2, LEU2, ura3, trp1-1, his3-11,15, can1-100, GAL, phi+, PSI+ snf5::KANMX* | S1B |
| UB18658 | MAT**a***, ADE2, leu2-3, ura3, trp1-1, his3-11,15, can1-100, GAL, psi+*<br>*snf5::KANMX*<br>*leu2::SNF5::LEU2* | S1B |
| UB18661 | MAT**a***, ADE2, leu2-3, ura3, trp1-1, his3-11,15, can1-100, GAL, psi+*<br>*snf5::KANMX*<br>*leu2::snf5-Q225X::LEU2* | S1B |
| UB41110 | MAT$\alpha$*, leu2-3,-112; his3-11,-15; trp1-1; ura3-1; ade1-4; can1-100, PSI+*<br>*YEp-Gal1p-SUP35NM-GFP:ura3* | S1G |
| UB41112 | MAT$\alpha$*, leu2-3,-112; his3-11,-15; trp1-1; ura3-1; ade1-4; can1-100, psi-*<br>*YEp-Gal1p-SUP35NM-GFP:ura3* | S1G |
| UB41897 (MAT**a**)<br>UB41898 (MAT$\alpha$)<br>UB41899 (MAT**a**) | *leu2-3,-112; his3-11,-15; trp1-1; ura3-1; ade1-4; can1-100, psi-, pin-, CAN1-370-TAA leu2::3V5-ade2-E64QQ::LEU2* | 2B; 2D; S2A |

| | | |
|---|---|---|
| UB41900 (MATα)<br>UB41901 (MAT**a**)<br>UB41902 (MATα) | *leu2-3,-112; his3-11,-15; trp1-1; ura3-1; ade1-4; can1-100, psi-, pin-, CAN1-370-TAA leu2::3V5-ade2-E64XQ::LEU2* | 2B; 2D; S2A |
| UB41891 (MATα)<br>UB41892 (MAT**a**)<br>UB41893 (MAT**a**) | *leu2-3,-112; his3-11,-15; trp1-1; ura3-1; ade1-4; can1-100, PSI+ leu2::3V5-ade2-E64QQ::LEU2* | 2B; 2D; S2A; S2B |
| UB41894 (MAT**a**)<br>UB41895 (MATα)<br>UB41896 (MATα) | *leu2-3,-112; his3-11,-15; trp1-1; ura3-1; ade1-4; can1-100, PSI+ leu2::3V5-ade2-E64XQ::LEU2* | 2B; 2D; S2A; S2B |
| UB48367 (MATα)<br>UB48368 (MAT**a**)<br>UB48369 (MATα) | *leu2-3,-112; his3-11,-15; trp1-1; ura3-1; ade1-4; can1-100, psi-, pin-, CAN1-370-TAA leu2::3V5-ade2-QE64Q::LEU2* | 2B; 2D; S2A |
| UB48364 (MAT**a**)<br>UB48365 (MATα)<br>UB48366 (MATα) | *leu2-3,-112; his3-11,-15; trp1-1; ura3-1; ade1-4; can1-100, psi-, pin-, CAN1-370-TAA leu2::3V5-ade2-QE64X::LEU2* | 2B; 2D; S2A |
| UB48340 (MAT**a**)<br>UB48341 (MATα)<br>UB48342 (MATα) | *leu2-3,-112; his3-11,-15; trp1-1; ura3-1; ade1-4; can1-100, PSI+ leu2::3V5-ade2-QE64Q::LEU2* | 2B; 2D; S2A |
| UB48337 (MATα)<br>UB48338 (MAT**a**)<br>UB48339 (MAT**a**) | *leu2-3,-112; his3-11,-15; trp1-1; ura3-1; ade1-4; can1-100, PSI+ leu2::3V5-ade2-QE64X::LEU2* | 2B; 2D; S2A |
| UB42459 (MATα)<br>UB42460 (MAT**a**)<br>UB42461 (MATα) | *leu2-3,-112; his3-11,-15; trp1-1; ura3-1; ade1-4; can1-100, psi-, pin-, CAN1-370-TAA leu2::3V5-ade2-QE64QQ-CAACAACAA::LEU2* | 2C; 2D; S2A |
| UB42462 (MAT**a**)<br>UB42463 (MATα)<br>UB42464 (MATα) | *leu2-3,-112; his3-11,-15; trp1-1; ura3-1; ade1-4; can1-100, psi-, pin-, CAN1-370-TAA leu2::3V5-ade2-QE64XQ-CAATAACAA::LEU2* | 2C; 2D; S2A |
| UB42453 (MAT**a**)<br>UB42454 (MATα)<br>UB42455 (MATα) | *leu2-3,-112; his3-11,-15; trp1-1; ura3-1; ade1-4; can1-100, PSI+ leu2::3V5-ade2-QE64QQ-CAACAACAA::LEU2* | 2C; 2D; S2A; S2B |
| UB42456 (MAT**a**)<br>UB42457 (MATα)<br>UB42458 (MATα) | *leu2-3,-112; his3-11,-15; trp1-1; ura3-1; ade1-4; can1-100, PSI+ leu2::3V5-ade2-QE64XQ-CAATAACAA::LEU2* | 2C; 2D; S2A; S2B |
| UB48373 (MATα)<br>UB48374 (MATα)<br>UB48375 (MAT**a**) | *leu2-3,-112; his3-11,-15; trp1-1; ura3-1; ade1-4; can1-100, psi-, pin-, CAN1-370-TAA leu2::3V5-ade2-QE64QQ-CAGCAGCAG::LEU2* | 2C; 2D; S2A |
| UB48606 (MAT**a**)<br>UB48607 (MATα)<br>UB48608 (MAT**a**) | *leu2-3,-112; his3-11,-15; trp1-1; ura3-1; ade1-4; can1-100, psi-, pin-, CAN1-370-TAA leu2::3V5-ade2-QE64XQ-CAGTAGCAG::LEU2* | 2C; 2D; S2A |
| UB48346 (MATα)<br>UB48347 (MATα)<br>UB48348 (MAT**a**) | *leu2-3,-112; his3-11,-15; trp1-1; ura3-1; ade1-4; can1-100, PSI+ leu2::3V5-ade2-QE64QQ-CAG***CAG***CAG::LEU2* | 2C; 2D; S2A; S2B |
| UB48603 (MATα)<br>UB48604 (MAT**a**)<br>UB48605 (MAT**a**) | *leu2-3,-112; his3-11,-15; trp1-1; ura3-1; ade1-4; can1-100, PSI+ leu2::3V5-ade2-QE64XQ-CAGTAGCAG::LEU2* | 2C; 2D; S2A; S2B |

| | | |
|---|---|---|
| UB49265 (MATa)<br>UB49266 (MATα)<br>UB49267 (MATa) | *leu2-3,-112; his3-11,-15; trp1-1; ura3-1; ade1-4; can1-100, psi-, pin-, CAN1-370-TAA leu2::3V5-ade2-QE64RQ-CAACGACAA::LEU2* | 2C; 2D; S2A |
| UB49315 (MATα)<br>UB49316 (MATα)<br>UB49317 (MATa) | *leu2-3,-112; his3-11,-15; trp1-1; ura3-1; ade1-4; can1-100, psi-, pin-, CAN1-370-TAA leu2::3V5-ade2-QE64XQ-CAATGACAA::LEU2* | 2C; 2D; S2A |
| UB49324 (MATa)<br>UB49325 (MATα)<br>UB49326 (MATa) | *leu2-3,-112; his3-11,-15; trp1-1; ura3-1; ade1-4; can1-100, PSI+, CAN1-370-TAA leu2::3V5-ade2-QE64RQ-CAACGACAA::LEU2* | 2C; 2D; S2A |
| UB49259 (MATα)<br>UB49260 (MATa)<br>UB49261 (MATα) | *leu2-3,-112; his3-11,-15; trp1-1; ura3-1; ade1-4; can1-100, PSI+, CAN1-370-TAA leu2::3V5-ade2-QE64XQ-CAATGACAA::LEU2* | 2C; 2D; S2A |
| UB49318 (MATα)<br>UB49319 (MATa)<br>UB49320 (MATα) | *leu2-3,-112; his3-11,-15; trp1-1; ura3-1; ade1-4; can1-100, psi-, pin-, CAN1-370-TAA leu2::3V5-ade2-QE64QQ-CAACAGCAA::LEU2* | 2C; 2D; S2A |
| UB49298(MATa)<br>UB49299 (MATα)<br>UB49300 (MATα) | *leu2-3,-112; his3-11,-15; trp1-1; ura3-1; ade1-4; can1-100, psi-, pin-, CAN1-370-TAA leu2::3V5-ade2-QE64XQ-CAATAGCAA::LEU2* | 2C; 2D; S2A |
| UB49262 (MATα)<br>UB49263 (MATα)<br>UB49264 (MATa) | *leu2-3,-112; his3-11,-15; trp1-1; ura3-1; ade1-4; can1-100, PSI+, CAN1-370-TAA leu2::3V5-ade2-QE64QQ-CAACAGCAA::LEU2* | 2C; 2D; S2A |
| UB49309(MATα)<br>UB49310 (MATa)<br>UB49311 (MATα) | *leu2-3,-112; his3-11,-15; trp1-1; ura3-1; ade1-4; can1-100, PSI+, CAN1-370-TAA leu2::3V5-ade2-QE64XQ-CAATAGCAA::LEU2* | 2C; 2D; S2A |
| UB49268 (MATα)<br>UB49269 (MATa)<br>UB49270 (MATα) | *leu2-3,-112; his3-11,-15; trp1-1; ura3-1; ade1-4; can1-100, psi-, pin-, CAN1-370-TAA leu2::3V5-ade2-QE64QQ-CAGCAGCAA::LEU2* | 2C; 2D; S2A |
| UB49301 (MATα)<br>UB49302 (MATα)<br>UB49303 (MATa) | *leu2-3,-112; his3-11,-15; trp1-1; ura3-1; ade1-4; can1-100, psi-, pin-, CAN1-370-TAA leu2::3V5-ade2-QE64XQ-CAGTAGCAA::LEU2* | 2C; 2D; S2A |
| UB49321 (MATα)<br>UB49322 (MATa)<br>UB49323 (MATa) | *leu2-3,-112; his3-11,-15; trp1-1; ura3-1; ade1-4; can1-100, PSI+, CAN1-370-TAA leu2::3V5-ade2-QE64QQ-CAGCAGCAA::LEU2* | 2C; 2D; S2A |
| UB49312 (MATα)<br>UB49313 (MATa)<br>UB49314 (MATα) | *leu2-3,-112; his3-11,-15; trp1-1; ura3-1; ade1-4; can1-100, PSI+, CAN1-370-TAA leu2::3V5-ade2-QE64XQ-CAGTAGCAA::LEU2* | 2C; 2D; S2A |
| UB48266 (MATa)<br>UB48267 (MATα) | *leu2-3,-112; his3-11,-15; trp1-1; ura3-1; ade1-4; can1-100, PSI+, CAN1-370-TAA leu2::3V5-ade2-E64QQ::LEU2 pdr5::KANMX* | S2C |
| UB48270 (MATa)<br>UB48271 (MATα) | *leu2-3,-112; his3-11,-15; trp1-1; ura3-1; ade1-4; can1-100, PSI+, CAN1-370-TAA leu2::3V5-ade2-QE64QQ::LEU2 pdr5::KANMX* | S2C |
| UB40396 (MATα)<br>UB40397 (MATa) | *leu2-3,-112; his3-11,-15; trp1-1; ura3-1; ade1-4; can1-100, psi-, pin-, CAN1-370-TAA* | 3C; 3D; S3A |

| | | |
|---|---|---|
| UB40398 (MAT*α*) | *leu2::3V5-CYC8::LEU2* | |
| UB40399 (MAT*α*)<br>UB40400 (MAT**a**)<br>UB40401 (MAT*α*) | *leu2-3,-112; his3-11,-15; trp1-1; ura3-1; ade1-4; can1-100, psi-, pin-, CAN1-370-TAA leu2::3V5-cyc8-Q648X::LEU2* | 3C; 3D; S3A |
| UB40402 (MAT*α*)<br>UB40403 (MAT**a**)<br>UB40404 (MAT*α*) | *leu2-3,-112; his3-11,-15; trp1-1; ura3-1; ade1-4; can1-100, PSI+, CAN1-370-TAA leu2::3V5-CYC8::LEU2* | 3C; 3D; S3A |
| UB40405 (MAT**a**)<br>UB40406 (MAT**a**)<br>UB40407 (MAT*α*) | *leu2-3,-112; his3-11,-15; trp1-1; ura3-1; ade1-4; can1-100, PSI+, CAN1-370-TAA leu2::3V5-cyc8-Q648X::LEU2* | 3C; 3D; S3A |
| UB37602 (MAT*α*)<br>UB37603 (MAT**a**)<br>UB37604 (MAT**a**) | *leu2-3,-112; his3-11,-15; trp1-1; ura3-1; ade1-4; can1-100, psi-, pin-, CAN1-370-TAA leu2::3V5-SPC25::LEU2* | 3C; 3D; S3B |
| UB37605 (MAT**a**)<br>UB37606 (MAT*α*)<br>UB37607 (MAT*α*) | *leu2-3,-112; his3-11,-15; trp1-1; ura3-1; ade1-4; can1-100, psi-, pin-, CAN1-370-TAA leu2::3V5-spc25-Q69X::LEU2* | 3C; 3D; S3B |
| UB37512 (MAT*α*)<br>UB37513 (MAT**a**)<br>UB37514 (MAT*α*) | *leu2-3,-112; his3-11,-15; trp1-1; ura3-1; ade1-4; can1-100, PSI+, CAN1-370-TAA leu2::3V5-SPC25::LEU2* | 3C; 3D; S3B |
| UB37515 (MAT*α*)<br>UB37516 (MAT**a**)<br>UB37517 (MAT**a**) | *leu2-3,-112; his3-11,-15; trp1-1; ura3-1; ade1-4; can1-100, PSI+, CAN1-370-TAA leu2::3V5-spc25-Q69X::LEU2* | 3C; 3D; S3B |
| UB37651 (MAT*α*)<br>UB37652 (MAT**a**)<br>UB37653 (MAT*α*) | *leu2-3,-112; his3-11,-15; trp1-1; ura3-1; ade1-4; can1-100, psi-, pin-, CAN1-370-TAA leu2::3V5-WHI3::LEU2* | 3C; 3D; S3C |
| UB37654 (MAT**a**)<br>UB37655 (MAT**a**)<br>UB37656 (MAT*α*) | *leu2-3,-112; his3-11,-15; trp1-1; ura3-1; ade1-4; can1-100, psi-, pin-, CAN1-370-TAA leu2::3V5-whi3-Q254X::LEU2* | 3C; 3D; S3C |
| UB37657 (MAT*α*)<br>UB37658 (MAT*α*)<br>UB37659 (MAT**a**) | *leu2-3,-112; his3-11,-15; trp1-1; ura3-1; ade1-4; can1-100, PSI+, CAN1-370-TAA leu2::3V5-WHI3::LEU2* | 3C; 3D; S3C |
| UB37660 (MAT**a**)<br>UB37661 (MAT*α*)<br>UB37662 (MAT**a**) | *leu2-3,-112; his3-11,-15; trp1-1; ura3-1; ade1-4; can1-100, PSI+, CAN1-370-TAA leu2::3V5-whi3-Q254X::LEU2* | 3C; 3D; S3C |
| UB45864 (MAT**a**)<br>UB45865 (MAT*α*) | *leu2-3,-112; his3-11,-15; trp1-1; ura3-1; ade1-4; can1-100, PSI+, CAN1-370-TAA cyc8::KANMX leu2::3V5-CYC8::LEU2* | 4B; S4F; S4G |
| UB45944 (MAT*α*)<br>UB45945 (MAT**a**) | *leu2-3,-112; his3-11,-15; trp1-1; ura3-1; ade1-4; can1-100, PSI+, CAN1-370-TAA cyc8::KANMX leu2::3V5-cyc8-Q648X::LEU2* | 4B; S4F; S4G |
| UB44142 (MAT**a**)<br>UB44143 (MAT*α*) | *leu2-3,-112; his3-11,-15; trp1-1; ura3-1; ade1-4; can1-100, PSI+, CAN1-370-TAA cyc8::KANMXcleu2::3V5-SNF5::LEU2* | 4C; S4F; S4H; S4I |
| UB45961 (MAT**a**)<br>UB45962 (MAT*α*) | *leu2-3,-112; his3-11,-15; trp1-1; ura3-1; ade1-4; can1-100, PSI+, CAN1-370-TAA* | 4C; S4F; S4H; S4I |

| | | |
|---|---|---|
| | *cyc8::KANMX leu2::3V5-snf5-Q225X::LEU2* | |
| UB45862 (MAT$\alpha$)<br>UB45863 (MAT**a**) | *leu2-3,-112; his3-11,-15; trp1-1; ura3-1; ade1-4; can1-100, psi-, pin-, CAN1-370-TAA snf5::KANMX leu2::3V5-SNF5::LEU2* | 4C; S4F; S4H; S4J |
| UB45646 (MAT$\alpha$)<br>UB45647 (MAT**a**) | *MAT$\alpha$/A, leu2-3,-112; his3-11,-15; trp1-1; ura3-1; ade1-4; can1-100, psi-, pin-, CAN1-370-TAA snf5::KANMX leu2::3V5-snf5-Q225X::LEU2* | 4C; S4F; S4H; S4J |
| UB41272 (MAT$\alpha$)<br>UB41273 (MAT**a**) | *leu2-3,-112; his3-11,-15; trp1-1; ura3-1; ade1-4; can1-100, psi-, pin-, CAN1-370-TAA UPF1 (WT cross control) leu2::3V5-SNF5::LEU2* | Table 1; Sup Data Table D-E |
| UB40561 (MAT**a**)<br>UB40562 (MAT$\alpha$) | *leu2-3,-112; his3-11,-15; trp1-1; ura3-1; ade1-4; can1-100, psi-, pin-, CAN1-370-TAA upf1Δ::HYG leu2::3V5-SNF5::LEU2* | Table 1; Sup Data Table D-E |
| UB41274 (MAT**a**)<br>UB41275 (MAT$\alpha$) | *leu2-3,-112; his3-11,-15; trp1-1; ura3-1; ade1-4; can1-100, psi-, pin-, CAN1-370-TAA UPF1 (WT cross control) leu2::3V5-snf5-Q225X::LEU2* | Table 1; Sup Data Table D-E |
| UB40558 (MAT$\alpha$)<br>UB40559 (MAT**a**) | *leu2-3,-112; his3-11,-15; trp1-1; ura3-1; ade1-4; can1-100, psi-, pin-, CAN1-370-TAA upf1Δ::HYG leu2::3V5-snf5-Q225X::LEU2* | Table 1; Sup Data Table D-E |
| UB41276 (MAT$\alpha$)<br>UB41277 (MAT**a**) | *leu2-3,-112; his3-11,-15; trp1-1; ura3-1; ade1-4; can1-100, PSI+, CAN1-370-TAA UPF1 (WT cross control) leu2::3V5-SNF5::LEU2* | Table 1; Sup Data Table D-E |
| UB41037 (MAT**a**)<br>UB41038 (MAT$\alpha$) | *leu2-3,-112; his3-11,-15; trp1-1; ura3-1; ade1-4; can1-100, PSI+, CAN1-370-TAA upf1Δ::HYG leu2::3V5-SNF5::LEU2* | Table 1; Sup Data Table D-E |
| UB41278 (MAT$\alpha$)<br>UB41279 (MAT**a**) | *leu2-3,-112; his3-11,-15; trp1-1; ura3-1; ade1-4; can1-100, PSI+, CAN1-370-TAA UPF1 (WT cross control) leu2::3V5-snf5-Q225X::LEU2* | Table 1; Sup Data Table D-E |
| UB40555 (MAT**a**)<br>UB40556 (MAT$\alpha$) | *leu2-3,-112; his3-11,-15; trp1-1; ura3-1; ade1-4; can1-100, PSI+, CAN1-370-TAA upf1Δ::HYG leu2::3V5-snf5-Q225X::LEU2* | Table 1; Sup Data Table D-E |
| UB48544 (MAT$\alpha$)<br>UB48545 (MAT**a**) | *leu2-3,-112; his3-11,-15; trp1-1; ura3-1; ade1-4; can1-100, psi-, pin-, CAN1-370-TAA EBS1 (WT cross control) leu2::3V5-SNF5::LEU2* | Table 1; Sup Data Table D-E |
| UB48542 (MAT**a**)<br>UB48543 (MAT$\alpha$) | *leu2-3,-112; his3-11,-15; trp1-1; ura3-1; ade1-4; can1-100, psi-, pin-, CAN1-370-TAA ebs1Δ::KANMX leu2::3V5-SNF5::LEU2* | Table 1; Sup Data Table D-E |
| UB48524 (MAT**a**)<br>UB48525 (MAT$\alpha$) | *leu2-3,-112; his3-11,-15; trp1-1; ura3-1; ade1-4; can1-100, psi-, pin-, CAN1-370-TAA EBS1 (WT cross control) leu2::3V5-snf5-Q225X::LEU2* | Table 1; Sup Data Table D-E |
| UB48522 (MAT**a**)<br>UB48523 (MAT$\alpha$) | *leu2-3,-112; his3-11,-15; trp1-1; ura3-1; ade1-4; can1-100, psi-, pin-, CAN1-370-TAA ebs1Δ::KANMX leu2::3V5-snf5-Q225X::LEU2* | Table 1; Sup Data Table D-E |

| | | |
|---|---|---|
| UB48520 (MAT$\alpha$)<br>UB48521 (MAT**a**) | *leu2-3,-112; his3-11,-15; trp1-1; ura3-1; ade1-4; can1-100, PSI+, CAN1-370-TAA*<br>*EBS1 (WT cross control) leu2::3V5-SNF5::LEU2* | Table 1; Sup Data Table D-E |
| UB48518 (MAT$\alpha$)<br>UB48519 (MAT**a**) | *leu2-3,-112; his3-11,-15; trp1-1; ura3-1; ade1-4; can1-100, PSI+, CAN1-370-TAA*<br>*ebs1Δ::KANMX leu2::3V5-SNF5::LEU2* | Table 1; Sup Data Table D-E |
| UB48548 (MAT$\alpha$)<br>UB48549 (MAT**a**) | *leu2-3,-112; his3-11,-15; trp1-1; ura3-1; ade1-4; can1-100, PSI+, CAN1-370-TAA*<br>*EBS1 (WT cross control) leu2:: snf5-Q225X::LEU2* | Table 1; Sup Data Table D-E |
| UB48546 (MAT**a**)<br>UB48547 (MAT$\alpha$) | *leu2-3,-112; his3-11,-15; trp1-1; ura3-1; ade1-4; can1-100, PSI+, CAN1-370-TAA*<br>*ebs1Δ::KANMX leu2:: snf5-Q225X::LEU2* | Table 1; Sup Data Table D-E |
| UB47488 (MAT**a**)<br>UB47489 (MAT$\alpha$) | *leu2-3,-112; his3-11,-15; trp1-1; ura3-1; ade1-4; can1-100, psi-, pin-, CAN1-370-TAA*<br>*DOM34 (WT cross control) leu2::3V5-SNF5::LEU2* | Table 1; Sup Data Table B |
| UB47486 (MAT**a**)<br>UB47487 (MAT$\alpha$) | *leu2-3,-112; his3-11,-15; trp1-1; ura3-1; ade1-4; can1-100, psi-, pin-, CAN1-370-TAA*<br>*dom34Δ::HYG leu2::3V5-SNF5::LEU2* | Table 1; Sup Data Table B |
| UB47492 (MAT**a**)<br>UB47493 (MAT$\alpha$) | *leu2-3,-112; his3-11,-15; trp1-1; ura3-1; ade1-4; can1-100, psi-, pin-, CAN1-370-TAA*<br>*DOM34 (WT cross control) leu2::3V5-snf5-Q225X::LEU2* | Table 1; Sup Data Table B |
| UB47490 (MAT**a**)<br>UB47491 (MAT$\alpha$) | *leu2-3,-112; his3-11,-15; trp1-1; ura3-1; ade1-4; can1-100, psi-, pin-, CAN1-370-TAA*<br>*dom34Δ::HYG leu2::3V5-snf5-Q225X::LEU2* | Table 1; Sup Data Table B |
| UB47496 (MAT$\alpha$)<br>UB47497 (MAT**a**) | *leu2-3,-112; his3-11,-15; trp1-1; ura3-1; ade1-4; can1-100, PSI+, CAN1-370-TAA DOM34 (WT cross control) leu2::3V5-SNF5::LEU2* | Table 1; Sup Data Table B |
| UB47494 (MAT$\alpha$)<br>UB47495 (MAT**a**) | *leu2-3,-112; his3-11,-15; trp1-1; ura3-1; ade1-4; can1-100, PSI+, CAN1-370-TAA*<br>*dom34Δ::HYG*<br>*leu2::3V5-SNF5::LEU2* | Table 1; Sup Data Table B |
| UB47500 (MAT**a**)<br>UB47501 (MAT$\alpha$) | *leu2-3,-112; his3-11,-15; trp1-1; ura3-1; ade1-4; can1-100, PSI+, CAN1-370-TAA*<br>*DOM34 (WT cross control) leu2::3V5-snf5-Q225X::LEU2* | Table 1; Sup Data Table B |
| UB47498 (MAT**a**)<br>UB47499 (MAT$\alpha$) | *leu2-3,-112; his3-11,-15; trp1-1; ura3-1; ade1-4; can1-100, PSI+, CAN1-370-TAA*<br>*dom34Δ::HYG leu2::3V5-snf5-Q225X::LEU2* | Table 1; Sup Data Table B |
| UB47534 (MAT**a**)<br>UB47535 (MAT$\alpha$) | *leu2-3,-112; his3-11,-15; trp1-1; ura3-1; ade1-4; can1-100, psi-, pin-, CAN1-370-TAA*<br>*ITT1 (WT cross control) leu2::3V5-SNF5::LEU2* | Table 1; Sup Data Table A |
| UB47532 (MAT**a**)<br>UB47533 (MAT$\alpha$) | *leu2-3,-112; his3-11,-15; trp1-1; ura3-1; ade1-4; can1-100, psi-, pin-, CAN1-370-TAA*<br>*itt1Δ::HYG leu2::3V5-SNF5::LEU2* | Table 1; Sup Data Table A |

| | | |
|---|---|---|
| UB47538 (MATα)<br>UB47539 (MAT**a**) | *leu2-3,-112; his3-11,-15; trp1-1; ura3-1; ade1-4; can1-100, psi-, pin-, CAN1-370-TAA ITT1 (WT cross control) leu2::3V5-snf5Q225X::LEU2* | Table 1; Sup Data Table A |
| UB47536 (MAT**a**)<br>UB47537 (MATα) | *leu2-3,-112; his3-11,-15; trp1-1; ura3-1; ade1-4; can1-100, psi-, pin-, CAN1-370-TAA itt1Δ::HYG leu2::3V5-snf5-Q225X::LEU2* | Table 1; Sup Data Table A |
| UB47542 (MAT**a**)<br>UB47543 (MATα) | *leu2-3,-112; his3-11,-15; trp1-1; ura3-1; ade1-4; can1-100, PSI+, CAN1-370-TAA ITT1 (WT cross control) leu2::3V5-SNF5::LEU2* | Table 1; Sup Data Table A |
| UB47540 (MAT**a**)<br>UB47541 (MATα) | *leu2-3,-112; his3-11,-15; trp1-1; ura3-1; ade1-4; can1-100, PSI+, CAN1-370-TAA itt1Δ::HYG leu2::3V5-SNF5::LEU2* | Table 1; Sup Data Table A |
| UB47546 (MATα)<br>UB47547 (MAT**a**) | *leu2-3,-112; his3-11,-15; trp1-1; ura3-1; ade1-4; can1-100, PSI+, CAN1-370-TAA ITT1 (WT control cross) leu2::3V5-snf5-Q225X::LEU2* | Table 1; Sup Data Table A |
| UB47544 (MAT**a**)<br>UB47545 (MATα) | *leu2-3,-112; his3-11,-15; trp1-1; ura3-1; ade1-4; can1-100, PSI+, CAN1-370-TAA itt1Δ::HYG leu2::3V5-snf5-Q225X::LEU2* | Table 1; Sup Data Table A |
| UB46615 (MAT**a**)<br>UB46616 (MATα) | *ade1-14 trp1-289 his3-Δ200 leu2- 3,112 ura3-52 OR leu2-3,-112; his3-11,-15; trp1-1; ura3-1; ade1-4 [psi-] rps30AΔ::natNT2 rps30BΔ YEplac112-RPS30B-FLAG-TRP1 leu2::3v5-SNF5::LEU2* | Table 1; Sup Data Table C |
| UB47558 (MAT**a**)<br>UB47559 (MATα) | *ade1-14 trp1-289 his3-Δ200 leu2- 3,112 ura3-52 OR leu2-3,-112; his3-11,-15; trp1-1; ura3-1; ade1-4 [psi-] rps30AΔ::natNT2 rps30BΔ YEplac112-RPS30B-FLAG-TRP1 leu2::3v5-snf5-Q267X::LEU2* | Table 1; Sup Data Table C |
| UB46617 (MATα)<br>UB46618 (MAT**a**) | *ade1-14 trp1-289 his3-Δ200 leu2- 3,112 ura3-52 OR leu2-3,-112; his3-11,-15; trp1-1; ura3-1; ade1-4 [PSI+] rps30AΔ::natNT2 rps30BΔ YEplac112-RPS30B-FLAG-TRP1 leu2::3v5-SNF5::LEU2* | Table 1; Sup Data Table C |
| UB47530 (MAT**a**)<br>UB47531 (MATα) | *ade1-14 trp1-289 his3-Δ200 leu2- 3,112 ura3-52 OR leu2-3,-112; his3-11,-15; trp1-1; ura3-1; ade1-4 [PSI+] rps30AΔ::natNT2 rps30BΔ YEplac112-RPS30B-FLAG-TRP1 leu2::3v5-snf5-Q267X::LEU2* | Table 1; Sup Data Table C |
| UB46451 (MATα)<br>UB46452 (MAT**a**) | *ade1-14 trp1-289 his3-Δ200 leu2- 3,112 ura3-52 OR leu2-3,-112; his3-11,-15; trp1-1; ura3-1; ade1-4 [psi-] rps30AΔ::natNT2 rps30BΔ YEplac112-RPS30B-R10A-FLAG-TRP1 leu2::3v5-SNF5::LEU2* | Table 1; Sup Data Table C |
| UB46455(MATα)<br>UB46456 (MAT**a**) | *ade1-14 trp1-289 his3-Δ200 leu2- 3,112 ura3-52 OR leu2-3,-112; his3-11,-15; trp1-1; ura3-1; ade1-4 [psi-] rps30AΔ::natNT2 rps30BΔ YEplac112-RPS30B-R10A-FLAG-TRP1 leu2::3v5-snf5Q267X::LEU2* | Table 1; Sup Data Table C |

| | | |
|---|---|---|
| UB46453 (MAT$\alpha$)<br>UB46454 (MAT**a**) | *ade1-14 trp1-289 his3-Δ200 leu2- 3,112 ura3-52 OR leu2-3,-112; his3-11,-15; trp1-1; ura3-1; ade1-4 [PSI+] rps30AΔ::natNT2 rps30BΔ YEplac112-RPS30B-R10A-FLAG-TRP1 leu2::3v5-SNF5::LEU2* | Table 1; Sup Data Table C |
| UB46458 (MAT$\alpha$)<br>UB46457 (MAT**a**) | *ade1-14 trp1-289 his3-Δ200 leu2- 3,112 ura3-52 OR leu2-3,-112; his3-11,-15; trp1-1; ura3-1; ade1-4 [PSI+] rps30AΔ::natNT2 rps30BΔ YEplac112-RPS30B-R10A-FLAG-TRP1 leu2::3v5-snf5-Q267X::LEU2* | Table 1; Sup Data Table C |
| UB46459 (MAT**a**)<br>UB46460 (MAT$\alpha$) | *leu2-3,-112; his3-11,-15; trp1-1; ura3-1; ade1-4; can1-100, PSI+, CAN1-370-TAA ura3::pRNR2-TetR-Tup1, pTetO7.1-TetR::URA3 leu2::TetOp-cyc8-Q648X-AAALELVDP-3V5::LEU2* | 5B-C; S5A-E |
| UB18045 (MAT$\alpha$)<br>UB18048 (MAT**a**) | *ade2-1, leu2-3, ura3, trp1-1, his3-11,15, can1-100, GAL, psi+ snf5::KANMX leu2::SNF5::LEU2* | 6B |
| UB18050 (MAT$\alpha$)<br>UB18052 (MAT**a**) | *ade2-1, leu2-3, ura3, trp1-1, his3-11,15, can1-100, GAL, psi+ snf5::KANMX leu2::snf5-Q225X::LEU2* | 6B |

**Supplementary Table 2. Primers used for qPCR and library barcoding.**

| Primer | Sequence |
|---|---|
| UB2598 (Act1 qPCR F) | gtaccaccatgttcccaggtatt |
| UB2599 (Act1 qPCR R) | agatggaccactttcgtcgt |
| UB9852 (3v5 universal qPCR F) | caaatcccttacttggtttgga |
| UB9853 (Snf5 qPCR R) | ggtaccctgcggctgattat |
| UB10816 (Ade1 qPCR R) | cgtcatatgcagagatacgatc |
| UB11481 (Ade2 qPCR R) | ccgtcttaatgttgagcctg |
| oCJC60 | caagcagaagacggcatacgagatattactcggtgactggagttcagacg |
| oCJC61 | caagcagaagacggcatacgagattccggagagtgactggagttcagacg |
| oCJC62 | caagcagaagacggcatacgagatcgctcattgtgactggagttcagacg |
| oCJC63 | caagcagaagacggcatacgagatgagattccgtgactggagttcagacg |
| oCJC64 | caagcagaagacggcatacgagatattcagaagtgactggagttcagacg |
| oCJC65 | caagcagaagacggcatacgagatgaattcgtgtgactggagttcagacg |
| oCJC66 | caagcagaagacggcatacgagatctgaagctgtgactggagttcagacg |
| oCJC67 | caagcagaagacggcatacgagattaatgcgcgtgactggagttcagacg |
| oCJC68 | caagcagaagacggcatacgagatcggctatggtgactggagttcagacg |
| oCJC69 | caagcagaagacggcatacgagattccgcgaagtgactggagttcagacg |
| oCJC70 | caagcagaagacggcatacgagattctcgcgcgtgactggagttcagacg |
| oCJC71 | caagcagaagacggcatacgagatagcgataggtgactggagttcagacg |
| oGBC72 | caagcagaagacggcatacgagatcacgatcagtgactggagttcagacg |
| oGBC73 | caagcagaagacggcatacgagatcgtatgtcgtgactggagttcagacg |
| oGBC74 | caagcagaagacggcatacgagatttaggcacgtgactggagttcagacg |
| oGBC75 | caagcagaagacggcatacgagattgtaccatgtgactggagttcagacg |
